# Promoter-sequence determinants and structural basis of primer-dependent transcription initiation in *Escherichia coli*

**DOI:** 10.1101/2021.04.06.438613

**Authors:** Kyle S. Skalenko, Lingting Li, Yuanchao Zhang, Irina O. Vvedenskaya, Jared T. Winkelman, Alexander Cope, Deanne M. Taylor, Premal Shah, Richard H. Ebright, Justin B. Kinney, Yu Zhang, Bryce E. Nickels

## Abstract

Chemical modifications of RNA 5′ ends enable “epitranscriptomic” regulation, influencing multiple aspects of RNA fate. In transcription initiation, a large inventory of substrates compete with nucleoside triphosphates (NTPs) for use as initiating entities, providing an *ab initio* mechanism for altering the RNA 5′ end. In *Escherichia coli* cells, RNAs with a 5′-end hydroxyl are generated by use of dinucleotide RNAs as primers for transcription initiation, “primer-dependent initiation.” Here we use massively systematic transcript end readout (“MASTER”) to detect and quantify RNA 5′ ends generated by primer-dependent initiation for ~4^10^ (~1,000,000) promoter sequences in *E. coli*. The results show primer-dependent initiation in *E. coli* involves any of the 16 possible dinucleotide primers and depends on promoter sequences in, upstream, and downstream of the primer binding site. The results yield a consensus sequence for primer-dependent initiation, Y_TSS-2_N_TSS-1_N_TSS_W_TSS+1_, where TSS is the transcription start site, N_TSS-1_N_TSS_ is the primer binding site, Y is pyrimidine, and W is A or T. Biochemical and structure-determination studies show that the base pair (nontemplate-strand base:template-strand base) immediately upstream of the primer binding site (Y:R_TSS-2_, where R is purine) exerts its effect through the base on the DNA template strand (R_TSS-2_) through inter-chain base stacking with the RNA primer. Results from analysis of a large set of natural, chromosomally-encoded *E*. *coli* promoters support the conclusions from MASTER. Our findings provide a mechanistic and structural description of how TSS-region sequence hard-codes not only the TSS position, but also the potential for epitranscriptomic regulation through primer-dependent transcription initiation.

## Introduction

In transcription initiation, the RNA polymerase (RNAP) holoenzyme binds promoter DNA by making sequence-specific interactions with core promoter elements and unwinds a turn of promoter DNA forming an RNAP-promoter open complex (RPo) containing a single-stranded “transcription bubble.” Next, RNAP selects a transcription start site (TSS) by placing the start-site nucleotide and the next nucleotide of the “template DNA strand” into the RNAP active-center product site (“P site”) and addition site (“A site”), respectively, and binding an initiating entity in the RNAP active-center P site (Figure 1A). RNAPs can initiate transcription using either a primer-independent or primer-dependent mechanism (1–10). In primer-independent initiation, the initiating entity (typically a nucleoside triphosphate, NTP) base pairs to the template-strand nucleotide in the RNAP active-center P site (TSS; Figure 1A). In primer-dependent transcription initiation, the 3′ nucleotide of a 2-, 3-, or 4-nucleotide RNA primer (di-, tri-, or tetranucleotide primer, respectively) base pairs to the template-strand nucleotide in the RNAP active-center P site, and the 5′ nucleotide of the primer base pairs to the template-strand nucleotide in the P-1, P-2, or P-3 site (TSS-1, TSS-2, and TSS-3, respectively; Figure 1A).

**Figure 1.**
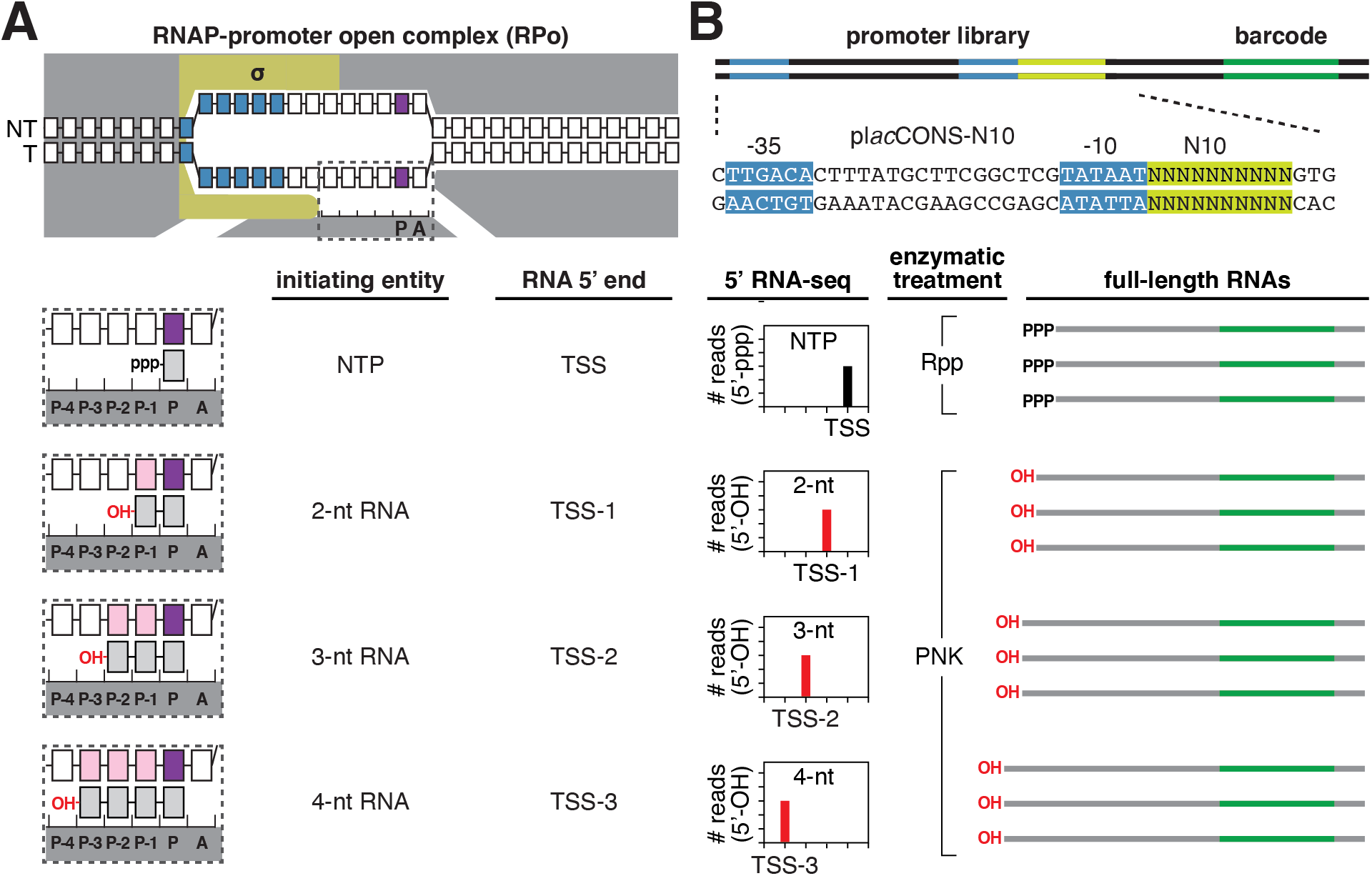
Use of massively systematic transcript end readout, “MASTER,” to monitor primer-independent and primer-dependent transcription initiation. **A.** Binding of initiating entities to DNA template-strand nucleotides in primer-independent and primer-dependent transcription initiation. Top: RNAP-promoter open complex (RPo). Bottom: Enlarged view of initiating entities bound to template-strand nucleotides in the RNAP active center. Dark gray, RNAP; yellow, σ; blue, −10-element nucleotides; purple, transcription start site (TSS) nucleotides; light gray, RNA nucleotides; pink, primer-binding nucleotides at positions TSS-1, TSS-2 or TSS-3; white boxes, DNA nucleotides; NT, nontemplate-strand nucleotides; T, template-strand nucleotides. P-3, P-2, P-1, and P, RNAP active-center initiating entity binding sites; A, RNAP active-center extending NTP binding site. Unwound transcription bubble in RPo indicated by raised and lowered nucleotides. **B.** Analysis of primer-independent and primer-dependent initiation using MASTER. Top: DNA fragment containing MASTER template library. Light green, randomized nucleotides in the promoter region; dark green, transcribed-region barcode. Bottom: 5′ RNA-seq analysis of RNA products generated from the promoter library by primer-independent, NTP-dependent initiation (Rpp treatment) and primer-dependent initiaion (PNK treatment).

In *Escherichia coli* cells, primer-dependent transcription initiation occurs during stationary-phase growth and modulates the expression of genes involved in biofilm formation (9–11). RNAs generated by primer-dependent initiation in *E. coli* contain a 5′-end hydroxyl (5′-OH), indicating that the primers incorporated at the RNA 5′ end also contain a 5′-OH (9). Available data suggests that most primer-dependent initiation in *E. coli* involves use of dinucleotide primers, most frequently UpA and GpG (9–11). However, direct evidence that dinucleotides serve as the predominant initiating entity in primer-dependent initiation has not been presented. In addition, apart from the sequences complementary to the primer, “the primer binding site,” promoter-sequence determinants for primer-dependent initiation have not been defined.

Here we adapt a massively parallel reporter assay to monitor primer-dependent initiation in *E. coli*. The results provide a complete inventory of the RNA 5′-end sequences generated by primer-dependent initiation in *E. coli* and define the critical promoter-sequence determinants for primer-dependent initiation. The results demonstrate that most, if not all, primer-dependent initiation in *E. coli* involves use of a dinucleotide as the initiating entity and identify a consensus sequence for primer-dependent initiation, Y_TSS-2_N_TSS-1_N_TSS_W_TSS+1_, where TSS is the transcription start site, N_TSS-1_N_TSS_ is the primer binding site, Y is pyrimidine, and W is A or T. We further demonstrate that sequence information at the position immediately upstream of the primer binding site resides exclusively in the template strand of the transcription bubble (R_TSS-2_, where R is purine). We report crystal structures of transcription initiation complexes containing dinucleotide primers that reveal the structural basis for a purine at the template-strand position immediately upstream of the primer binding site (R_TSS-2_): namely, more extensive, and likely more energetically favorable, base stacking between the template-strand base and the primer 5′ base.

## Results

### Use of massively systematic transcript end readout, “MASTER,” to monitor primer-dependent initiation in *E. coli*

To define, comprehensively, the promoter-sequence determinants for primer-dependent initiation in *E. coli*, we modified a massively parallel reporter assay previously developed in our lab termed massively systematic transcript end readout, “MASTER” (12, 13), in order to detect both primer-independent and primer-dependent initiation, to differentiate between primer-independent and primer-dependent initiation, and to define primer lengths in primer-dependent initiation (Figure 1B).

MASTER involves construction of a promoter library that contains up to 4^11^ (~4,000,000) barcoded sequences, production of RNA transcripts from the promoter library, and analysis of RNA barcodes and RNA 5′ ends using high-throughput sequencing (5′ RNA-seq) to define, for each RNA product, the template that produced the RNA and the sequence of the RNA 5′ end (Figure 1B) (12–14). The 5′ RNA-seq procedure used in MASTER relies on ligation of single-stranded oligonucleotide adaptors to RNAs containing a 5′-end monophosphate (5′-p) (13). In previous work, we treated RNAs with RNA 5′ Pyrophosphohydrolase (Rpp), which converts a 5′-end triphosphate (5′-ppp) to a 5′-p; this procedure specifically enables detection of the 5′-ppp bearing RNAs generated by primer-independent initiation (12, 14–16). Here, we treated RNAs, in parallel, with Rpp to detect RNAs generated by primer-independent initiation and with Polynucleotide Kinase (PNK), which converts a 5′-OH to a 5′-p, to detect RNAs generated by primer-dependent initiation (Figure 1B). By comparing the results from Rpp and PNK reactions we quantify, for each promoter sequence in the library, the relative efficiencies of primer-independent and primer-dependent initiation, and primer lengths for primer-dependent initiation.

In the present work, we used a MASTER template library containing 4^10^ (~1,000,000) sequence variants at the positions 1-10 base pairs (bp) downstream of the −10 element of a consensus σ^70^-dependent promoter (p*lac*CONS-N10; Figure 1B). The randomized segment of p*lac*CONS-N10 contains the full range of TSS positions for *E. coli* RNAP observed in previous work (i.e., TSS positions located 6, 7, 8, 9, and 10 bp downstream of the promoter −10 element; Figure 2A). We introduced the p*lac*CONS-N10 library into *E. coli*, grew cells to stationary phase (the phase in which primer-dependent initiation has been observed in previous work; 9), isolated total cellular RNA, and analyzed RNAs generated from each promoter sequence in the library by 5′ RNA-seq. The results provide complete inventories of RNA 5′ ends generated by primer-independent initiation and primer-dependent initiation in stationary-phase *E. coli* cells.

**Figure 2.**
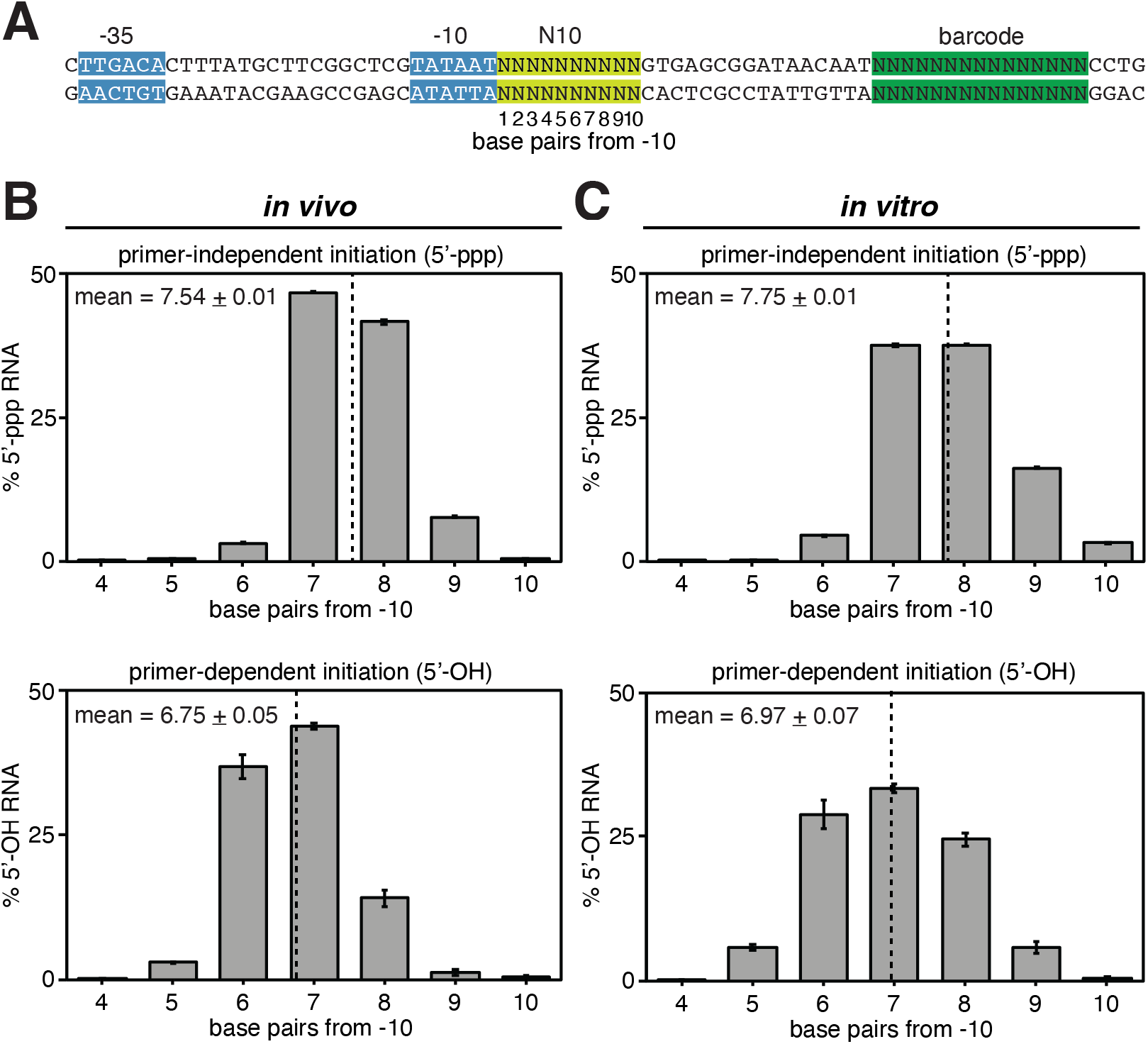
Distributions of 5′-end sequences for RNAs generated in primer-independent initiation and primer-dependent initiation *in vivo* and *in vitro*. **A.** *lac*CONS-N10 library. Base pairs in the N10 region are numbered based on their position relative to the promoter −10 element. Colors as in Figure 1B. **B-C.** RNA 5′-end distribution histograms (mean ± SD, N = 3) for RNAs generated by primer-independent initiation (top) or primer-dependent initiation (bottom) in stationary-phase *E. coli* cells (panel B) or *in vitro*, with the dinucleotide primer UpA (panel C). Dashed line indicates the mean 5′-end position (mean ± SD, N = 3).

### Primer-dependent initiation: 5′-end positions

Our results define distributions of 5′-end positions of the 5′-ppp RNAs generated by primer-independent initiation and the 5′-OH RNAs generated by primer-dependent initiation for transcription in stationary-phase *E. coli* cells (Figure 2B).

The distributions of 5′-end positions for primer-independent initiation show 5′-end positions (TSS positions) ranging from 6-10 bp downstream of the promoter −10 element, with a mean 5′-end position ~7.5 bp downstream of the promoter −10 element (Figure 2B, top). The range, the mean, and the distribution shape closely match those previously observed for primer-independent initiation for cells in exponential phase (12).

The distributions of 5′-end positions for primer-dependent initiation show 5′-end positions ranging from 5-9 bp downstream of the promoter −10 element, with a mean 5′-end position ~6.8 bp downstream of the promoter −10 element (Figure 2B, bottom). The range, the mean, and the distribution shape closely match those for primer-independent initiation, but with a ~1 bp upstream shift.

### Primer-dependent initiation: primer lengths

Comparison of the 5′-end distributions for primer-independent initiation (Figure 2B, top) vs. primer-dependent initiation (Figure 2B, bottom) indicates that, across all promoter sequences in the library, the 5′-end positions of RNAs generated by primer-independent initiation (mean position 7.54 ± 0.01 bp downstream of −10 element) and RNAs generated by primer-dependent initiation (mean position 6.75 ± 0.05 bp downstream of −10 element) differ by ~1 bp (0.71 ± 0.06 bp). Following the logic of Figure 1, based on the observed difference of almost exactly 1 bp in mean 5′-end position for primer-independent initiation vs. primer-dependent initiation, we infer that primer length in primer-dependent initiation in stationary-phase *E. coli* cells is almost always 2 nt. Computational modeling, using the distributions in Figure 2B, indicates that no more than ~2.5% of the observed primer-dependent initiation could involve primer lengths greater than 2-nt (Figure S1A). Consistent with these inferences, comparison of distributions of RNA 5′-end positions for primer-independent initiation *in vitro* vs. primer-dependent initiation *in vitro* with the dinucleotide primer UpA shows essentially the same ~1 bp upstream shift in distribution range, mode, and mean (Figures 2C, S1B).

### Primer-dependent initiation: primer sequences

We next measured yields of 5′-OH RNAs generated by primer-dependent initiation with each of the 16 possible dinucleotide primers (Figure 3A). The results show that primer-dependent initiation occurs with all 16 dinucleotide primers. Highest levels of primer-dependent initiation are observed with the dinucleotide primers UpA and GpG, which account for ~27% and ~17%, respectively, of 5′-OH RNAs generated across all promoters in the library (Figure 3A, left). The other 14 dinucleotide primers each account for ~1% to ~8% of 5′-OH RNAs generated across all promoters in the library. Qualitatively similar results are obtained analyzing RNA products from promoters where the primer binding site is at positions 5-6, 6-7, 7-8, 8-9, and 9-10 bp downstream of the promoter −10 element (Figure 3A, right). The demonstration that primer-dependent initiation occurs with all 16 dinucleotides *in vivo* is new to this work, as is the demonstration that primer-dependent initiation occurs at the full range of TSS positions observed for primer-independent initiation *in vivo* (i.e., TSS positions located 6, 7, 8, 9, and 10 bp downstream of the promoter −10 element). The observation that UpA and GpG are preferentially used as primers *in vivo* is consistent with results of prior work (9, 10).

**Figure 3.**
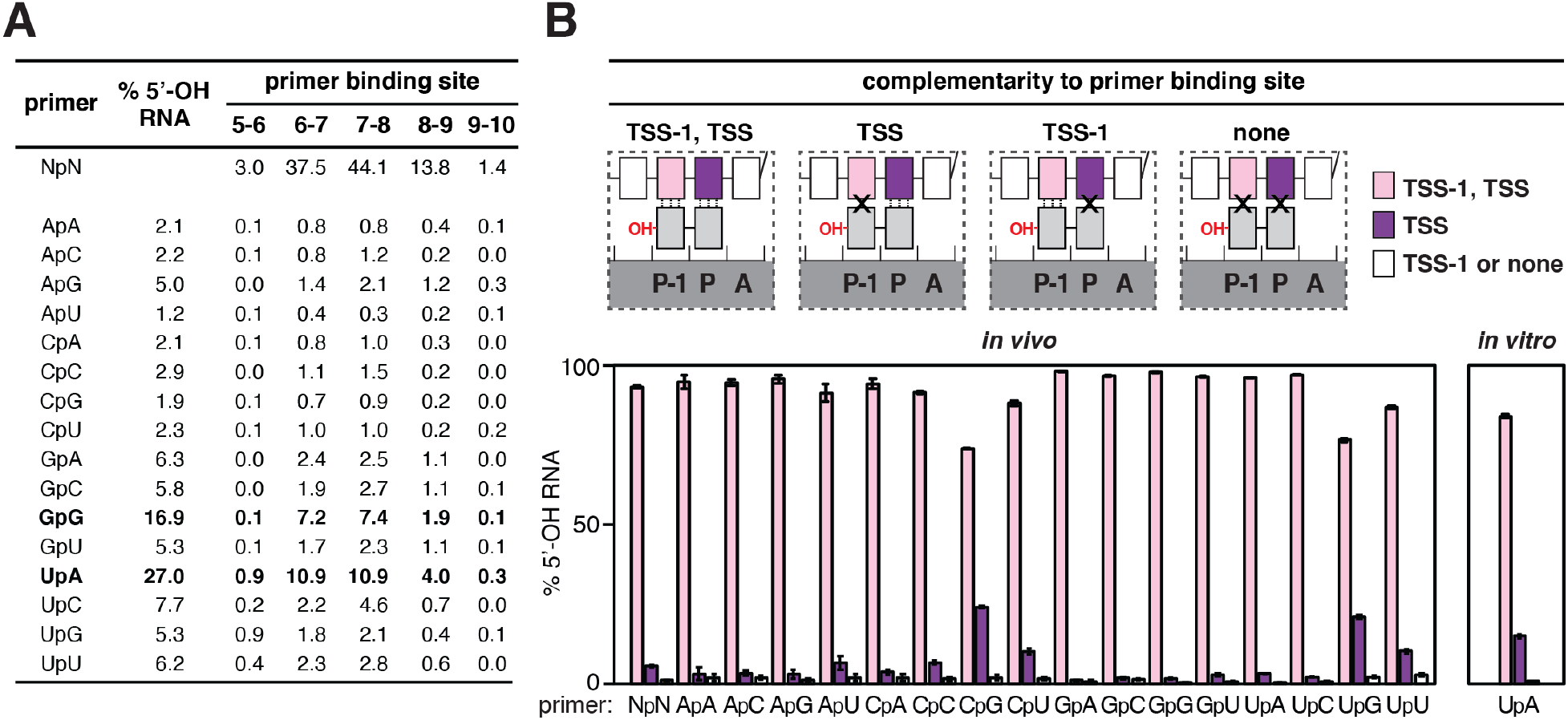
Promoter-sequence dependence of primer-dependent initiation: primer binding site. **A.** Relative usage of dinucleotides in primer-dependent initiation in stationary-phase *E. coli* cells. Values represent the percentage of total 5′-OH RNAs generated using each of the 16 dinucleotide primers (mean, N = 3). Bold, dinucleotides preferentially used as primers. **B.** Complementarity between the primer binding site and dinucleotide in primer-dependent initiation. Top: primer-dependent initiation involving template-strand complementarity to both 5′ and 3′ nucleotides of primer (TSS-1, TSS), template-strand complementarity to only 3′ nucleotide of primer (TSS), template-strand complementarity to only 5′ nucleotide of primer (TSS-1), or no template-strand complementarity to primer (none). Three vertical lines, complementarity; X, non-complementarity. Other symbols and colors as in Figure 1. Bottom: percentage of primer-dependent initiation involving complementarity to both 5′ and 3′ nucleotides of primer (TSS-1, TSS; pink), complementarity to only 3′ nucleotide of primer (TSS; purple), or template-strand complementarity to only 5′ nucleotide of primer or no template-strand complementarity to primer (TSS-1 or none; white) in stationary-phase *E. coli* cells (left) or *in vitro*, with the dinucleotide primer UpA (right) (mean ± SD, N = 3).

### Primer-dependent initiation: promoter-sequence dependence, primer binding site

Analysis of the results for primer-dependent initiation, separately considering RNA products with 5′ ends complementary to the template and RNA products with 5′ ends non-complementary to the template, shows that the overwhelming majority of primer-dependent initiation in stationary-phase *E. coli* cells occurs at primer binding sites that have perfect template-strand complementarity to the 5′ and 3′ nucleotides of the dinucleotide primer (93.3 ± 0.4%; Figure 3B, bottom). This is true, across the entire promoter library, for each of the 16 possible dinucleotide primer sequences (73.9 ± 0.2% to 98.1 ± 0.01% of primer binding sites with perfect complementarity; Figure 3B, bottom), and for each of the major primer binding-site positions (Figure S2). Most of the limited non-complementarity observed involves the 5′ nucleotide of the dinucleotide primers CpG, UpG, CpU and UpU (24.2 ± 0.4%, 21.1 ± 0.6%, 10.3 ± 0.6%, and 10.1 ± 0.9%, respectively; Figures 3B, S2). Consistent with these results, analysis of the same promoter library *in vitro*, assessing primer-dependent initiation with the dinucleotide primer UpA, shows that the overwhelming majority of primer-dependent initiation likewise occurs at primer binding sites that have perfect template-strand complementarity to the 5′ and 3′ nucleotides of the dinucleotide primer for each of the major primer binding-site positions (84.1 ± 0.6%; Figures 3B, bottom right, S3). *In vitro* transcription experiments using heteroduplex templates (templates having non-complementary transcription-bubble nontemplate-strand and template-strand sequences) and the dinucleotide primer UpA show that the strong preference for perfect template-strand complementarity to the 5′ and 3′ nucleotides of the dinucleotide primer reflects a requirement for Watson-Crick base pairing of template-strand nucleotides at positions TSS-1 and TSS with the 5′ and 3′ nucleotides of the dinucleotide RNA primer, respectively (Figure S4).

We conclude that primer-dependent initiation with a dinucleotide primer almost always involves a primer binding site having perfect template-strand complementarity to, and therefore able to engage in Watson-Crick base pairing with, the dinucleotide primer. This result is not completely unexpected. However, this point has not been demonstrated previously *in vivo*, and prior work *in vitro*, with tetranucleotide primers (17), had indicated that perfect template-strand complementarity to the primer may not be necessary for primer-dependent initiation with longer primers.

### Primer-dependent initiation: promoter-sequence dependence, sequences flanking the primer binding site

The observed yields of 5′-OH RNA products from primer-dependent initiation in stationary-phase *E. coli* cells strongly correlate with the promoter sequences flanking the primer binding site (Figures 4, S5). Levels of primer-dependent initiation depend on the identities of base pairs up to 7 bases upstream of the primer binding site and up to 3 bases downstream of the primer binding site. The base pair (nontemplate-strand base:template-strand base) at the position immediately upstream of the primer binding site, position TSS-2, makes the largest contribution to the sequence dependence of primer-dependent initiation. Levels of primer-dependent initiation are higher for promoters with a nontemplate-strand pyrimidine (C or T) and template-strand purine (A or G) at position TSS-2 (Figure 4, left). A strong preference for a Y:R base pair at position TSS-2 is observed at each of the major TSS positions (i.e., the positions 6, 7, and 8 bp downstream of the promoter −10 element; Figure 4, left), The results further show that the base pair at the position immediately downstream of the primer binding site, position TSS+1, makes the second largest contribution to the sequence dependence of primer-dependent initiation; levels of primer-dependent initiation are higher for promoters with a T:A or A:T base pair at position TSS+1 (Figure 4, left). A strong preference for T:A or A:T at position TSS+1 is observed when the TSS is 6 bp downstream of the promoter −10 element, and a weaker preference is observed when the TSS is 7 or 8 bp downstream of the promoter −10 element (Figure 4, left). Base pairs at positions 3, 4, 5, 6, and 7 bp upstream of the primer binding site (TSS-3, TSS-4, TSS-5, TSS-6, and TSS-7) and at positions 2 and 3 bp downstream of the primer binding site (TSS+2 and TSS+3), make small, but significant, contributions to levels of primer-dependent initiation (Figure 4, left). The results define a global consensus sequence for primer-dependent initiation: Y_TSS-2_N_TSS-1_N_TSS_W_TSS+1_ (Y:R_TSS-2_N:N_TSS-1_N:N_TSS_W:W_TSS+1_), where TSS is the transcription start site, N_TSS-1_N_TSS_ is the primer binding site, Y is pyrimidine, and W is A or T. The same or essentially the same consensus is observed for all 16 primer sequences and for each major primer binding-site position (Figure S5). Analysis of the same promoter library *in vitro*, assessing primer-dependent initiation with the dinucleotide primer UpA, yields the same consensus sequence, Y_TSS-2_N_TSS-1_N_TSS_W_TSS+1_, and does so for each primer binding-site position (Figures 4, right, and S6).

**Figure 4.**
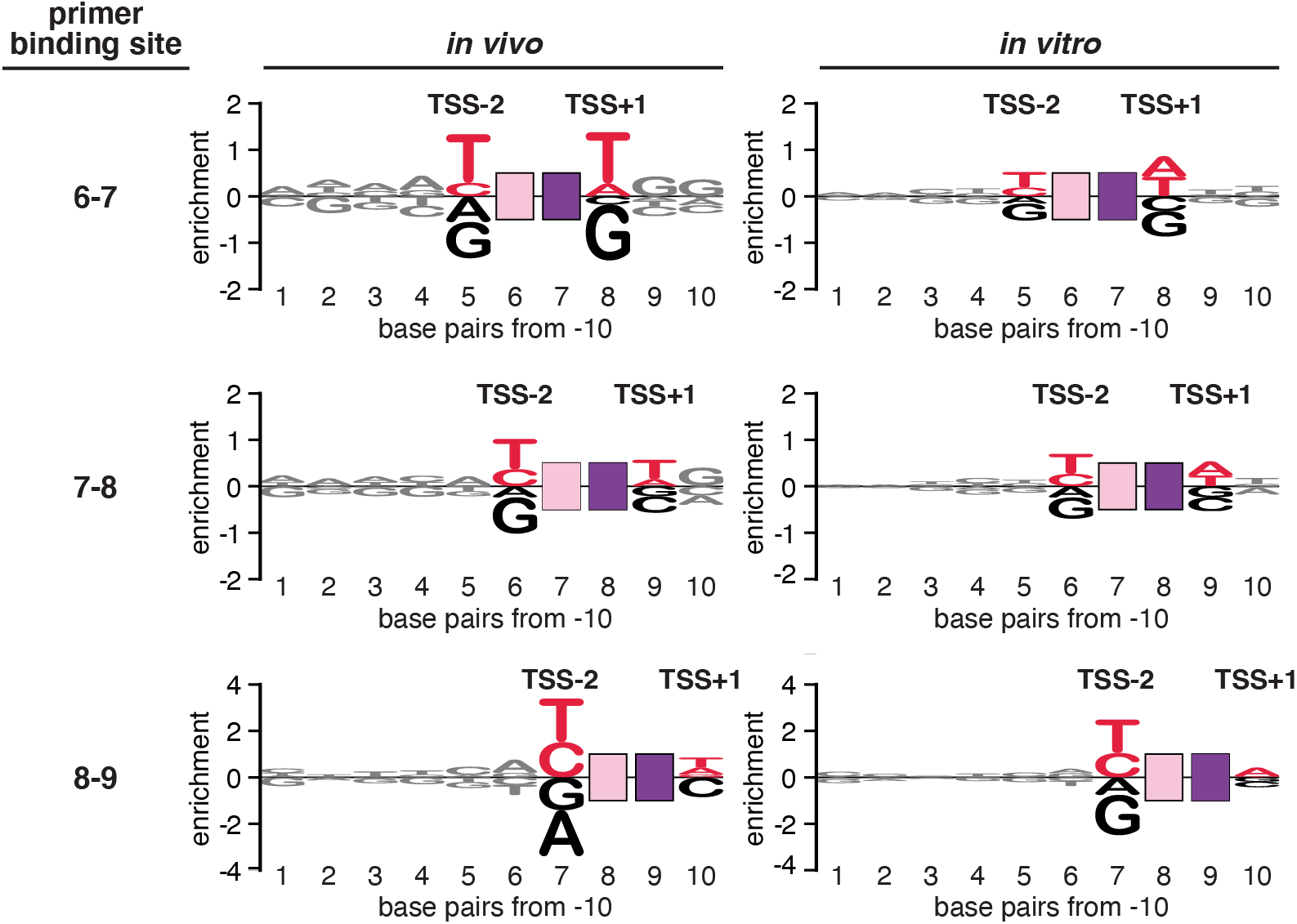
Promoter-sequence dependence of primer-dependent initiation: sequences flanking the primer binding site. Sequence logo (35) for primer-dependent initiation at TSS positions 7, 8, and 9 (corresponding to primer binding sites 6-7, 7-8, and 8-9, respectively) in stationary-phase *E. coli* cells (left) or *in vitro*, with the dinucleotide primer UpA (right). The height of each base “X” at each position “Y” represents the log_2_ average of the % 5′-OH RNAs computed across sequences containing nontemplate-strand X at position Y. Red, consensus nucleotides; black, non-consensus nucleotides. Other symbols and colors as in Figure 1.

*In vitro* transcription experiments assessing competition between primer-dependent transcription initiation with UpA and primer-independent transcription initiation with ATP show that primer-dependent initiation is ~60 times more efficient than primer-independent initiation at a promoter conforming to the consensus sequence (T_TSS-2_T_TSS-1_A_TSS_T_TSS+1_; Figure 5A), but is only ~10 times more efficient at a promoter not having the consensus sequence (G_TSS-2_T_TSS-1_A_TSS_T_TSS+1_; Figure 5A).

**Figure 5.**
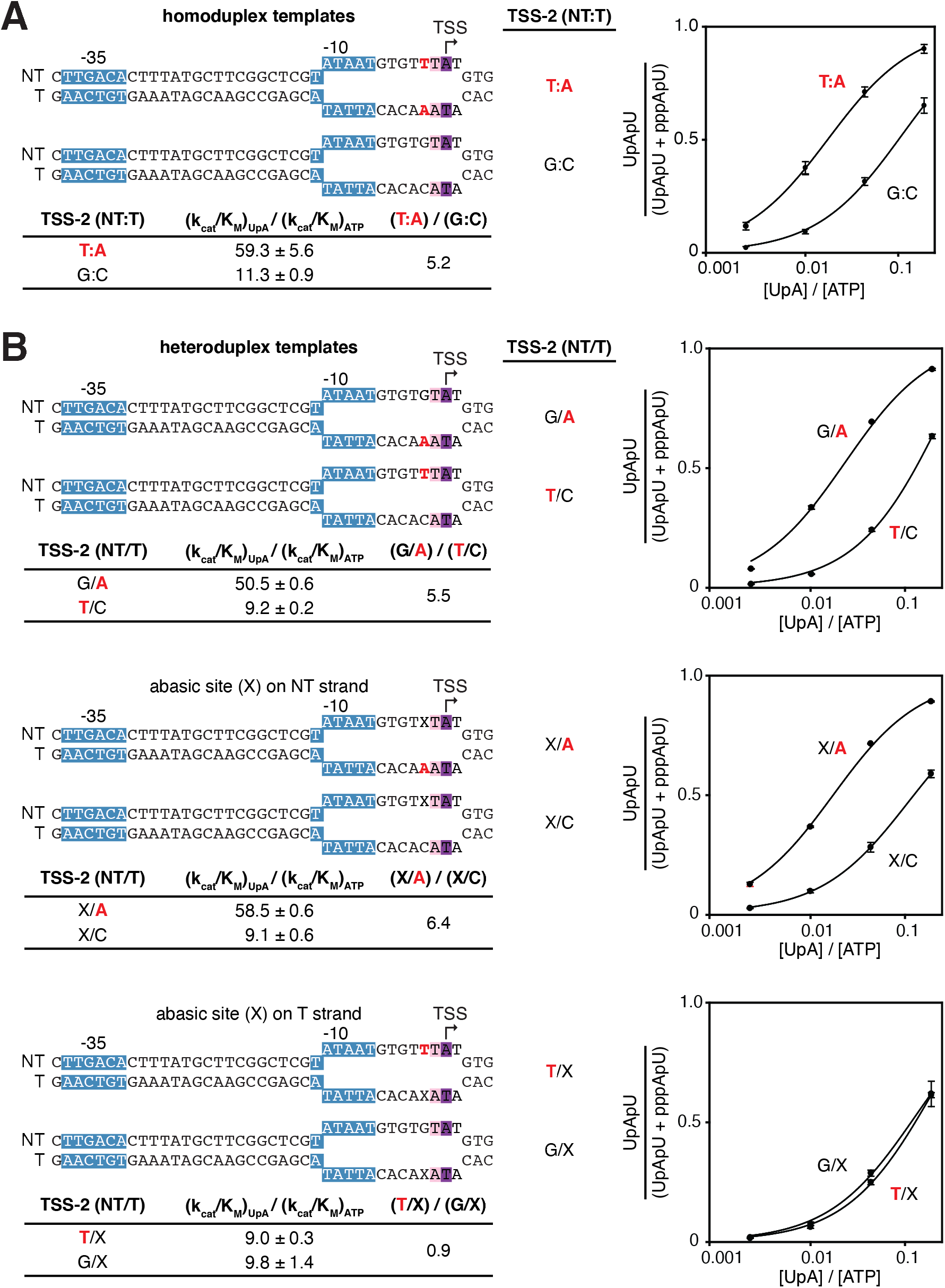
Promoter-sequence dependence of primer-dependent initiation *in vitro*: position TSS-2. **A.** Relative efficiencies of primer-dependent initiation vs. primer-independent initiation depends on promoter sequence at position TSS-2. Top left: homoduplex DNA templates containing consensus or non-consensus nucleotides for primer-dependent initiation at position TSS-2. Bottom left: relative efficiencies of primer-dependent initiation with UpA vs. primer-independent initiation with ATP [(k_cat_/K_M_)_UpA_ / (k_cat_/K_M_)_ATP_], and ratio of (k_cat_/K_M_)_UpA_ / (k_cat_/K_M_)_ATP_ for the indicated templates. Right: dependence of primer-dependent initiation on [UpA] / [ATP] ratio (mean ± SD, N = 3). Red, consensus nucleotides at position TSS-2. Unwound transcription bubble in RPo indicated by raised and lowered nucleotides. Other symbols and colors as in Figure 1. **B.** The template DNA strand carries sequence information at position TSS-2. Top left: DNA templates containing mismatches at position TSS-2. Templates contain a consensus nucleotide at position TSS-2 on only the nontemplate strand (T/C_TSS-2_ and T/X_TSS-2_), only the template strand (G/A_TSS-2_ and X/A_TSS-2_), or neither strand (X/C_TSS-2_ and G/X_TSS-2_). Bottom left: relative efficiencies and efficiency ratios for the indicated heteroduplex templates. Right: dependence of primer-dependent initiation on [UpA] / [ATP] ratio (mean ± SD, N = 3). Red, consensus nucleotides at position TSS-2. Unwound transcription bubble in RPo indicated by raised and lowered nucleotides.

*In vitro* transcription experiments using heteroduplex templates and the dinucleotide primer UpA show that the sequence information responsible for the preference for Y:R at TSS-2 resides exclusively in the DNA template strand (Figure 5B). Thus, in experiments with heteroduplex templates, primer-dependent initiation is reduced by replacement of the consensus nucleotide by a non-consensus nucleotide or an abasic site on the DNA template strand, but is not reduced by replacement of the consensus nucleotide by a non-consensus nucleotide or an abasic site on the DNA nontemplate strand (Figure 5B).

We conclude that primer-dependent initiation, *in vivo* and *in vitro*, depends not only on the sequence of the primer binding site, but also on flanking sequences, with the preferred sequence being Y_TSS-2_N_TSS-1_N_TSS_W_TSS+1_ (Y:R_TSS-2_N:N_TSS-1_N:N_TSS_W:W_TSS+1_).

### Primer-dependent initiation: chromosomal promoters

To assess whether the sequence preferences observed in the MASTER analysis apply also to natural promoters, we quantified primer-dependent initiation in stationary-phase *E. coli* cells at each of 93 promoters that use UpA as primer (Table S1, Figure 6A). The results show the same sequence preferences at positions TSS-2 and TSS+1 observed in the MASTER analysis are observed in chromosomally-encoded promoters (Figure 6A).

**Figure 6.**
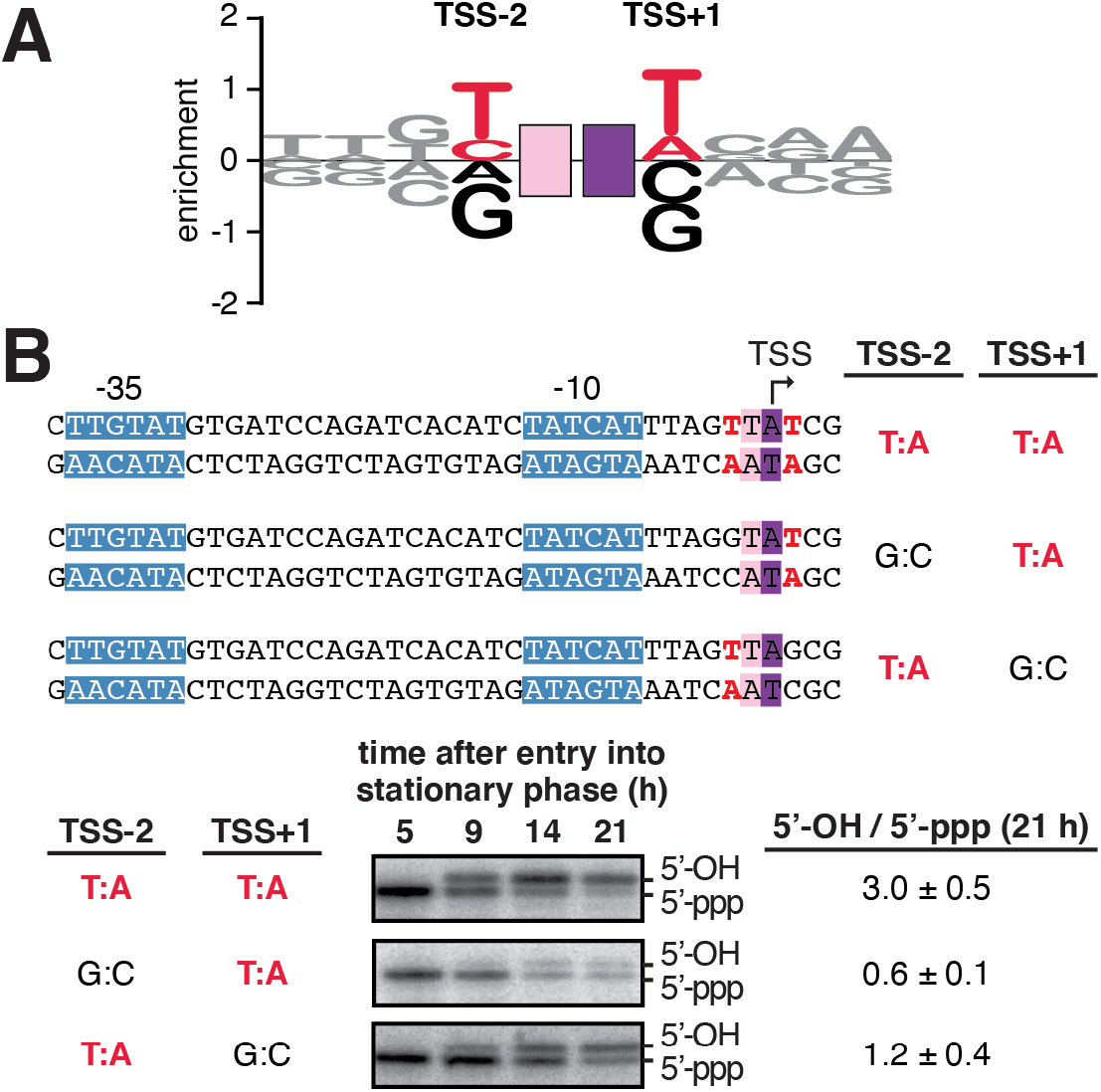
Promoter-sequence dependence of primer-dependent initiation: chromosomal promoters. **A.** Sequence logo (35) for primer-dependent initiation at TSS positions 7, 8, and 9 (corresponding to primer binding sites 6-7, 7-8, and 8-9, respectively) in stationary-phase *E. coli* cells for 93 natural, chromosomally-encoded promoters that use UpA as a primer. The height of each base “X” at each position “Y” represents the log_2_ average of the % 5′-OH RNAs computed across sequences containing nontemplate-strand X at position Y. Red, consensus nucleotides; black, non-consensus nucleotides. Other symbols and colors as in Figure 1. **B.** Promoter-sequence dependence of primer-dependent initiation at the *E. coli bhsA* promoter. Top: sequences of DNA templates containing wild-type and mutant derivatives of *bhsA* promoter. Bottom: primer extension analysis of 5′-end lengths of *bhsA* RNAs. In primer-dependent initiation with a dinucleotide primer, the RNA product acquires one additional nucleotide at the RNA 5′ end (Figure 1). Gel shows radiolabeled cDNA products derived from primer-independent initiation (5′-ppp) and primer-dependent initiation (5′-OH) in stationary-phase *E. coli* cells. Bottom right: ratios of primer-dependent initiation vs. primer-independent initiation (mean ± SD, N = 4).

To assess directly the functional significance of the sequence preferences observed in the MASTER analysis and natural promoter analysis, we constructed mutations at positions TSS-2 and TSS+1 of a natural promoter that uses UpA as primer (Figure 6B, top) and assessed effects on function in stationary-phase *E. coli* cells (Figure 6B, bottom). We observed that, at position TSS-2, the consensus base pair T:A is preferred over the non-consensus base pair G:C by a factor of ~5 (Figure 6B, bottom), and, at position TSS+1, the consensus base pair T:A is preferred over the non-consensus base pair G:C by a factor of ~2.5 (Figure 6B, bottom). We conclude that the sequence dependence for primer-dependent initiation defined using MASTER is also observed in natural, chromosomally-encoded *E. coli* promoters.

### Primer-dependent initiation: structural basis of promoter sequence-dependence

To determine the structural basis of the preference for a template-strand purine nucleotide at position TSS-2 (R_TSS-2_) in primer-dependent initiation, we determined crystal structures of transcription initiation complexes containing a template-strand purine nucleotide at position TSS-2 (Figure 7). We first prepared crystals of *Thermus thermophilus* RPo using synthetic nucleotide scaffolds containing a template-strand purine nucleotide, A, at position TSS-2, and containing a template-strand primer binding site for either the dinucleotide primer used most frequently in primer-dependent initiation *in vivo*, UpA (9, 10; Figure 3A), or the dinucleotide primer used second most frequently in primer-dependent initiation *in vivo*, GpG (9, 10; Figure 3A). We next soaked the crystals either with UpA and CMPcPP or with GpG and CMPcPP, to yield crystals of *T. thermophilus* RPo in complex with a dinucleotide primer and a non-reactive analog of an extending NTP. We then collected X-ray diffraction data, solved structures, and refined structures, obtaining structures of RPo[A_TSS-2_A_TSS-1_T_TSS_]-UpA-CMPcPP at 2.8 Å resolution and RPo[A_TSS-2_C_TSS-1_C_TSS_]-GpG-CMPcPP at 2.9 Å resolution (Table S2, Figure 7).

**Figure 7.**
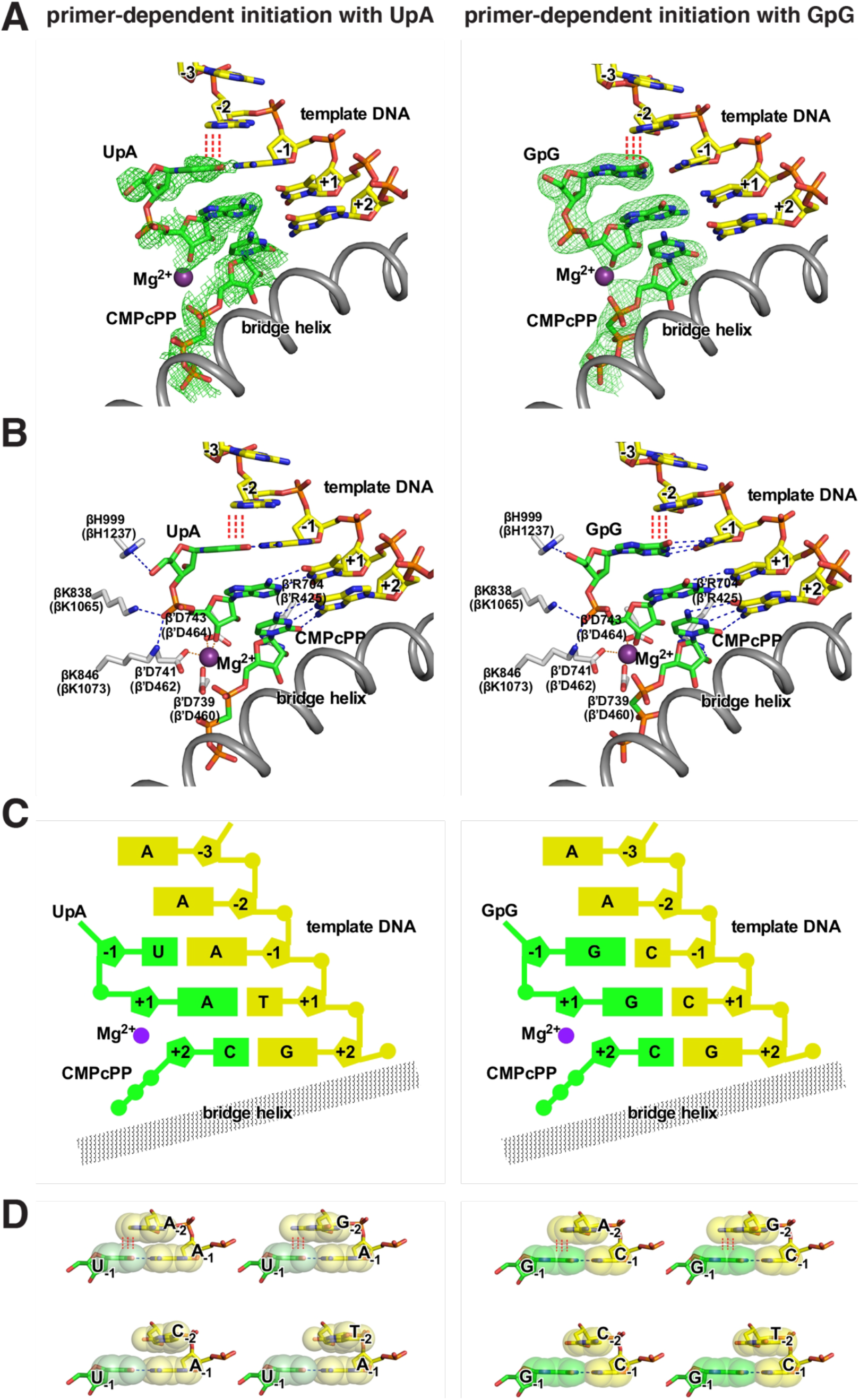
Structural basis of promoter-sequence dependence of primer-dependent initiation at position TSS-2. Crystal structures of *T. thermophilus* RPo[ATSS-2ATSS-1TTSS]-UpA-CMPcPP (left) and *T. thermophilus* RPo[A_TSS-2_C_TSS-1_C_TSS_]-GpG-CMPcPP (right). **A.** Experimental electron density (contoured at 2.5σ; green mesh) and atomic model for DNA template strand (yellow, red, blue, and orange for C, O, N, and P atoms), dinucleotide primer and CMPcPP (green, red, blue, and orange for C, O, N, and P atoms), RNAP active-center catalytic Mg^2+^(I) (violet sphere), and RNAP bridge helix (gray ribbon). **B.** Contacts of RNAP residues (gray, red, and blue for C, O, and N atoms) with primer and RNAP active-center catalytic Mg^2+^(I). RNAP residues are numbered both as in *T. thermophilus* RNAP and as in *E. coli* RNAP (in parentheses). **C.** Schematic summary of structures. Template-strand DNA (yellow); primer and CMPcPP (green); RNAP bridge helix (gray); RNAP active-center catalytic Mg^2+^(I) (violet). **D.** Structural basis of promoter-sequence dependence at position TSS-2. Extensive inter-chain base stacking of template-strand purine, A or G, with 5′ nucleotide of primer (upper row; red vertical dashed lines), and limited inter-chain base stacking of template-strand pyrimidine, C or T, and 5′ nucleotide of primer (lower row). The inter-chain base-stacking patterns of template-strand A with primers UpA and GpG are as observed in structures of RPo[A_TSS-2_A_TSS-1_T_TSS_]-UpA-CMPcPP and RPo[A_TSS-2_C_TSS-1_C_TSS_]-GpG-CMPcPP (panels A-C); the inter-chain base-stacking pattern of template-strand T with primer GpG is as observed in structure of RPo[T_TSS-2_C_TSS-1_C_TSS_]-GpG-CMPcPP (Figure S7); the other inter-chain base-stacking patterns are modeled by analogy. Base atoms are shown as van der Waals surfaces. Colors are as in panel A.

For both RPo[A_TSS-2_A_TSS-1_T_TSS_]-UpA-CMPcPP and RPo[A_TSS-2_C_TSS-1_C_TSS_]-GpG-CMPcPP, experimental electron density maps show unambiguous density for the dinucleotide primer in the RNAP active-center P-1 and P sites and for CMPcPP in the RNAP active-center A site (Figure 7B). The dinucleotide primers make extensive interactions with RNAP, template-strand DNA, and CMPcPP. For each dinucleotide primer, the primer phosphate, makes the same interactions with RNAP (residues βH1237, βK1065 and βK1073; residues numbered as in *E. coli* RNAP) and the RNAP active-center catalytic Mg^2+^ as the primer phosphate in a previously reported structure of *T. thermophilus* RPo-GpA (18; Figure 7A-C). For each dinucleotide primer, the primer bases make Watson-Crick H-bonds with template-strand nucleotides at TSS-1 and TSS, and make an intra-chain stacking interaction with the base of CMPcPP. Crucially, for each dinucleotide primer, the 5′ base of the primer makes an inter-chain stacking interaction with the base of the template-strand purine nucleotide at position TSS-2 (Figure 7). Structural modeling indicates that this inter-chain base-stacking interaction can occur only when the template-strand nucleotide at position TSS-2 is a purine (Figure 7D). Structural modeling further indicates that this inter-chain base-stacking interaction should stabilize the binding of the primer and CMPcPP to template-strand DNA, thereby facilitating primer-dependent initiation. Consistent with this structural modeling, a crystal structure of a complex obtained by soaking crystals of *T. thermophilus* RPo containing a template-strand pyrimidine nucleotide, T, at position TSS-2 with GpG and CMPcPP (RPo[T_TSS-2_C_TSS-1_C_TSS_]-GpG; 3.4 Å resolution; Table S2) do not show significant inter-chain base-stacking with the template-strand at position TSS-2, and do not show binding of CMPcPP to template-strand DNA (Figure S7). Taken together, the crystal structures in Figures 7 and S7 define the structural basis of the preference for purine vs. pyrimidine at template-strand position TSS-2 in primer-dependent transcription initiation: namely, inter-chain base stacking between the primer 5′ base and a purine at template-strand position TSS-2 facilitates binding of the primer and the extending NTP.

## Discussion

### Promoter-sequence dependence of primer-dependent initiation: mechanism and structural basis

Our biochemical results show that primer-dependent initiation in stationary-phase *E. coli* almost always involves a dinucleotide primer (Figures 2, S1), can involve any of the 16 possible dinucleotide primers (Figure 3A), almost always involves a primer binding site complementary to both the 5′ and 3′ nucleotides of the dinucleotide primer (Figures 3B, S2–S4), depends on promoter sequences flanking the primer binding site (Figures 4–6, S5–S6), and exhibits the consensus sequence Y_TSS-2_N_TSS-1_N_TSS_W_TSS+1_ (Y:R_TSS-2_N:N_TSS-1_N:N_TSS_W:W_TSS+1_; Figures 4–6, S5–S6), wherein the sequence information at positions TSS-2, TSS-1, and TSS is contained exclusively within the promoter template strand (Figures 5, S4).

Our structural results show that the sequence preference for purine at template-strand position TSS-2 is a consequence of inter-chain base stacking of a purine at template-strand position TSS-2 with the 5′ nucleotide of a dinucleotide primer (Figures 7, S7). The structural basis of the preference for purine vs. pyrimidine at template-strand position TSS-2 in primer-dependent initiation is analogous to--almost identical to--the previously described structural basis of the preference for purine vs. pyrimidine at template-strand position TSS-1 in primer-independent initiation (19, 20). In the former case, inter-chain base stacking between the primer 5′ base in the RNAP active-center P-1 site and a purine at template-strand position TSS-2 facilitates binding of the primer and an extending NTP. In the latter case, inter-chain base-stacking between the initiating NTP in the RNAP active-center P site and a purine at template-strand position TSS-1 facilitates binding of the initiating NTP and an extending NTP.

### Promoter sequences upstream of the TSS modulate the chemical nature of the RNA 5′ end

Chemical modifications of the RNA 5′ end provide a layer of “epitranscriptomic” regulation, influencing multiple aspects of RNA fate, including stability, processing, localization, and translation efficiency (21–24). Primer-dependent initiation provides one mechanism to alter the RNA 5′ end during transcription initiation. In primer-dependent initiation with a dinucleotide primer, the RNA product acquires a 5′ hydroxyl and acquires one additional nucleotide at the RNA 5′ end (Figure 1). In an analogous manner, NCIN-dependent initiation--where an NCIN is a non-canonical initiating nucleotide--provides another mechanism to alter the RNA 5′ end during transcription initiation (25, 26). In NCIN-dependent initiation, the RNA product acquires an NCIN at the RNA 5′ end. NCIN-dependent initiation has been shown to occur with the oxidized and reduced forms of nicotinamide adenine dinucleotide (NAD), dephospho-coenzyme A (dpCoA), flavin adenine dinucleotide (FAD), uridine diphosphate N-acetylglucosamine (UDP-GlcNAc), and dinucleoside tetraphosphates (Np4Ns) (25, 27–30)

All three modes of transcription initiation--primer-dependent, NCIN-dependent, and NTP-dependent--exhibit promoter-sequence dependence (12, 16, 25, 30; Figures 3–6, S2–S6). All three modes of transcription initiation exhibit promoter consensus sequences that include base pairs upstream of the initiating-entity binding site (12, 16, 25, 30; Figures 4–6, S5–S6). Crucially, the promoter consensus sequences upstream of the initiating-entity binding site for primer-dependent, NCIN-dependent (NAD, dpCoA, and Np4N), and NTP-dependent transcription initiation all are different: Y:R_TSS-2_, R:Y_TSS-1_, and Y:R_TSS-1_, respectively (12, 16, 25, 30; Figures 4–6, S5–S6). It follows that the sequence of the promoter TSS region hard-codes not only the TSS position, but also the relative efficiencies of, and potentials for epitranscriptomic regulation through, primer-dependent, NCIN-dependent, and NTP-dependent transcription initiation.

## Materials and Methods

### Proteins

*E. coli* RNAP core enzyme used in transcription experiments was prepared from *E. coli* strain NiCo21(DE3) (New England Biolabs, NEB) transformed with plasmid pIA900 (31) using culture and induction procedures, immobilized-metal-ion affinity chromatography on Ni-NTA agarose, and affinity chromatography on Heparin HP as described in (32). *E. coli* σ^70^ was prepared from *E. coli* strain NiCo21 (DE3) transformed with plasmid pσ^70^-His using culture and induction procedures, immobilized-metal-ion affinity chromatography on Ni-NTA agarose, and anion-exchange chromatography on Mono Q as described in (33). 10x RNAP holoenzyme was formed by mixing 0.5 μM RNAP core and 2.5 μM σ^70^ in 1x reaction buffer (40 mM Tris HCl, pH 7.5; 10 mM MgCl_2_; 150 mM KCl; 0.01% Triton X-100; and 1 mM DTT).

5’ RNA polyphosphatase (Rpp) and T4 Polynucleotide Kinase (PNK) were purchased from Epicentre and NEB, respectively.

### Oligonucleotides

Oligodeoxyribonucleotides (Table S3) were purchased from Integrated DNA Technologies (IDT) and were purified with standard desalting purification. UpA and GpG were purchased from Trilink Biotechnologies. NTPs (ATP, GTP, CTP, and UTP) were purchased from GE Healthcare Life Sciences.

Homoduplex and heteroduplex templates used in single-template *in vitro* transcription assays were generated by mixing 1.1 μM nontemplate-strand oligo with 1 μM template-strand oligo in 10 mM Tris (pH 8.0). Mixtures were heated to 90°C for 10 min and slowly cooled to 40°C (0.1°C / second) using a Dyad PCR machine (Bio-Rad).

### Plasmids

Plasmid pBEN516 (9) contains sequences from −100 to +15 of the *bhsA* promoter fused to the tR’ terminator inserted between the HindIII and SalI sites of pACYC184 (NEB). Mutant derivatives of pBEN516 containing a G:C base pair at position TSS-2 (pKS494) or a G:C base pair at position +2 (pKS497) were generated using site-directed mutagenesis. Plasmid pPSV38 (9) contains a pBR322 origin, a gentamycin resistance gene (*aaC1*), and *lacI*q. Plasmid library pMASTER-*lac*CONS-N10 has been previously described (12).

### Analysis of primer-dependent initiation by MASTER (Figures 2–4, S1–S3, S5–S6)

#### Primer-dependent initiation in vitro: transcription reaction conditions

A linear DNA fragment containing p*lac*CONS-N10 generated as described in (34) was used as template for *in vitro* transcription assays. Transcription reactions (total volume = 100 μl) were performed by mixing 10 nM of template DNA with 50 nM RNAP holoenzyme in *E. coli* RNA polymerase reaction buffer (NEB) (40 mM Tris HCl, pH 7.5; 10 mM MgCl_2_; 150 mM KCl; 0.01% Triton X-100; 1 mM DTT), 0.1 mg/ml BSA (NEB), and 40 U murine RNase inhibitor (NEB). Reactions were incubated at 37°C for 15 min to form open complexes. A single round of transcription was initiated by addition of 1000 μM ATP, 1000 μM CTP, 1000 μM UTP, 1000 μM GTP, UpA (40 μM, 160 μM, or 640 μM), and 0.1 mg/ml heparin (Sigma Aldrich). Reactions were incubated at 37°C for 15 min and stopped by addition of 0.5 M EDTA (pH 8) to a final concentration of 50 mM. (For each replicate, two 100 μl transcription reactions were performed separately and combined after addition of EDTA.) Nucleic acids were recovered by ethanol precipitation, reconstituted in 25 μl of nuclease-free water, mixed with 25 μl of 2x RNA loading dye (95% formamide; 25 mM EDTA; 0.025% SDS; 0.025% xylene cyanol; 0.025% bromophenol blue), and separated by electrophoresis on 10% 7M urea slab gels (Invitrogen) equilibrated and run in 1x TBE. The gel was stained with SYBR Gold nucleic acid gel stain (Invitrogen), bands visualized on a UV transilluminator, and RNA products ~150 nt in length were excised from the gel. The excised gel slice was crushed, 300 μl of 0.3 M NaCl in 1x TE buffer was added, and the mixture was incubated at 70°C for 10 min. Eluted RNAs were collected using a Spin-X column (Corning). After the first elution, the crushed gel fragments were collected and the elution procedure was repeated, nucleic acids were collected, pooled with the first elution, isolated by isopropanol precipitation, and resuspended in 25.5 μl of RNase-free water (Invitrogen). Reactions were performed in triplicate.

#### Primer-dependent initiation in stationary-phase E. coli cells: cell growth

Three independent 25 ml cell cultures of *E. coli* MG1655 cells (gift of A. Hochschild, Harvard Medical School) containing p*lac*CONS-N10 and pPSV38 were grown in LB media (Millipore) containing chloramphenicol (25 μg/ml), gentamicin (10 μg/ml), and IPTG (1 mM) in a 125 ml DeLong flask (Bellco Glass) shaken at 220 RPM at 37°C until late stationary phase (~21 hours after entry into stationary phase; final OD600 ~3.5). 2 ml aliquots of cell suspensions were placed in 2 ml tubes and cells were collected by centrifugation (1 min; 21,000 x *g*; 20°C). Supernatants were removed and cells stored at −80°C.

### Primer-dependent initiation in stationary-phase E. coli cells: RNA isolation

RNA was isolated from frozen cell pellets as described in (12). Cell pellets were resuspended in 600 μl of TRI Reagent solution (Molecular Research Center), incubated at 70°C for 10 min, and centrifuged (10 min; 21,000 x *g*; 4°C) to remove insoluble material. The supernatant was transferred to a fresh tube, ethanol was added to a final concentration of 60.5%, and the mixture was applied to a Direct-zol spin column (Zymo Research). DNase I (Zymo Research) treatment was performed on-column according to the manufacturer’s recommendations. RNA was eluted from the column using nuclease-free water heated to 70°C (3 x 30 μl elutions; total volume of eluate = 90 μl). RNA was treated with 2 U TURBO DNase (Invitrogen) at 37°C for 1 h, samples were extracted with acid phenol:chloroform (Ambion), RNA was recovered by ethanol precipitation and resuspended in RNase-free water. A MICROBExpress Kit (Invitrogen) was used to remove rRNAs from ~36 μg of recovered RNA, rRNA-depleted RNA was isolated by ethanol precipitation and resuspended in 40 μl of RNase-free water.

### Enzymatic treatment of RNA products

For RNAs isolated from *E. coli*, 3 μg of rRNA-depleted RNA was used in each reaction. RNAs isolated from *in vitro* transcription reactions were split into four equal portions and used in each reaction.

Rpp treatment (total reaction volume = 30 μl): RNA products were mixed with 20 U Rpp and 40 U RNaseOUT (Invitrogen) in 1x Rpp reaction buffer (50 mM HEPES-KOH, pH 7.5; 100 mM NaCl; 1 mM EDTA; 0.1% BME; and 0.01% Triton X-100) and incubated at 37°C for 1 hr. Reactions were extracted with acid phenol:chloroform, RNA was recovered by ethanol precipitation, and resuspended in 10.5 μl RNase-free water.

PNK treatment (total reaction volume = 50 μl): RNA products were mixed with 20 U PNK, 40 U RNaseOUT, and 1 mM ATP (NEB) in 1x PNK reaction buffer (70 mM Tris-HCl pH 7.6, 10 mM MgCl_2_, 5 mM DTT) and incubated at 37°C for 1 hr. Processed RNAs were recovered using Qiagen’s RNeasy MinElute kit (following the manufacturer’s recommendations with the exception that RNAs were eluted from the column using 200 μl nuclease-free water heated to 70°C). RNA was recovered by ethanol precipitation and resuspended in 10.5 μl RNase-free water.

Rpp and PNK treatment: RNA products were mixed with 20 U PNK, 40 U RNaseOUT, and 1 mM ATP in 1x PNK reaction buffer (total reaction volume = 50 μl) and incubated at 37°C for 1 hr. Processed RNAs were recovered using Qiagen’s RNeasy MinElute kit (following the manufacturer’s recommendations with the exception that RNAs were eluted from the column using 25 μL nuclease-free water heated to 70°C). Recovered RNA products were mixed with 20 U Rpp and 40 U RNaseOUT in 1x Rpp reaction buffer (total reaction volume = 30 μl) and incubated at 37°C for 1 hr. Reactions were extracted with acid phenol:chloroform, RNA was recovered by ethanol precipitation, and resuspended in 10.5 μl RNase-free water.

“mock” PNK treatment (total reaction volume = 50 μl): RNA products were mixed with 40 U RNaseOUT and 1 mM ATP in 1x PNK reaction buffer and incubated at 37°C for 1 hr. Reactions were extracted with acid phenol:chloroform, RNA was recovered by ethanol precipitation, and resuspended in 10.5 μl RNase-free water.

“mock” Rpp treatment (total reaction volume = 30 μl): RNA products were mixed with 40 U RNaseOUT in 1x Rpp reaction buffer (total reaction volume = 30 μl) and incubated at 37°C for 1 hr. Reactions were extracted with acid phenol:chloroform, RNA was recovered by ethanol precipitation, and resuspended in 10.5 μl RNase-free water.

#### 5’-adaptor ligation

To enable quantitative comparisons between samples, we performed the 5’-adaptor ligation step using barcoded 5’-adaptor oligonucleotides as described in (16). For RNA products isolated from stationary-phase *E. coli* cells, oligo i105 was used for RNAs processed by Rpp, oligo i106 was used for RNAs processed with PNK, oligo i107 was used for RNAs processed with both Rpp and PNK, and oligo i108 was used for unprocessed RNAs (mock PNK treated). For RNA products isolated from *in vitro* reactions, oligo i105 was used for RNAs processed by Rpp, oligo i106 was used for unprocessed RNAs (mock Rpp treated); oligo i107 was used for RNAs processed by PNK, and oligo i108 was used for unprocessed RNAs (mock PNK treated).

Processed RNA products isolated from stationary-phase *E. coli* cells (in 10.5 μl of nuclease-free water) were combined with 1 mM ATP (NEB), 40 U RNaseOUT, 1x T4 RNA ligase buffer (NEB), and 10 U T4 RNA ligase 1 (NEB) and 1 μM of 5’ adaptor oligo (total reaction volume = 20 μl), and incubated at 37°C for 2 h. Reactions were then supplemented with 1x T4 RNA ligase buffer, 1 mM ATP, PEG 8000 (10% final), 5U T4 RNA ligase 1, and 20 U RNaseOUT (total reaction volume = 30 μl) and further incubated at 16°C for 16 h. Processed RNA products isolated from *in vitro* reactions (in 10.5 μl of nuclease-free water) were combined with PEG 8000 (10% final concentration), 1 mM ATP, 40 U RNaseOUT, 1x T4 RNA ligase buffer, 10 U T4 RNA ligase 1, and 1 μM of 5’ adaptor oligo (total reaction volume = 30 μl), and incubated at 16°C for 16 h.

Ligation reactions were stopped by addition of 30 μl of 2x RNA loading dye and heated at 95°C for 5 min. For each replicate, the 4 ligation reactions were combined, and separated by electrophoresis on 10% 7M urea slab gels (equilibrated and run in 1x TBE). Gels were incubated with SYBR Gold nucleic acid gel stain, and bands were visualized with UV transillumination. For RNAs isolated from stationary-phase *E. coli* cells, products migrating above the 5’-adapter oligo were isolated from the gel (procedures as above; recovered in 50 μl of nuclease-free water). For RNAs generated *in vitro*, products ~150 nt in length were recovered from the gel (procedures as above; recovered in 16 μl of nuclease-free water). 5’-adaptor-ligated RNAs were used for analysis of primer-dependent initiation from p*lac*CONS-10 (this section) and for analysis of primer-dependent initiation from natural, chromosomally-encoded promoters (next section).

#### First strand cDNA synthesis

For RNAs isolated from stationary-phase *E. coli* cells, 25 μl of 5’-adaptor-ligated RNAs were mixed with 1.5 μl s128A oligonucleotide (3 μM) and 3.5 μl nuclease-free water. The 30 μl mixture was incubated at 65°C for 5 min, cooled to 4°C, and combined with 20 μl of a solution containing 10 μl of 5x First-Strand buffer (Invitrogen), 2.5 μl of 10 mM dNTP mix (NEB), 2.5 μl of 100 mM DTT (Invitrogen), 2.5 μl 40 U/μl RNaseOUT, and 2.5 μl 100 U/μl SuperScript III Reverse Transcriptase (Invitrogen), for a final reaction volume of 50 μl. Reactions were incubated at 25°C for 5 min, 55°C for 60 min, 70°C for 15 min, then kept at 25°C. Next, 5.4 μl of 1M NaOH was added, reactions were incubated at 95°C for 5 min, and kept at 10°C, 4.5 μl 1.2M HCl was added, followed by 60 μl of 2x RNA loading dye.

For RNAs isolated from *in vitro* reactions, 16 μl of 5’-adaptor-ligated RNAs were mixed with 0.5 μl s128A oligonucleotide (1.5 μM). The 16.5 μl mixture was incubated at 65°C for 5 min, cooled to 4°C, and combined with 13.5 μl of a solution containing 6 μl of 5x First-Strand buffer, 1.5 μl of 10 mM dNTP mix, 1.5 μl of 100 mM DTT, 1 μl 40 U/μl RNaseOUT, 1.5 μl 100 U/μl SuperScript III Reverse Transcriptase, and 2 μl of nuclease-free water, for a final reaction volume of 30 μl. Reactions were incubated at 25°C for 5 minutes, 55°C for 60 minutes, 70°C for 15 min, then cooled to 25°C. Next, 10 U of RNase H (NEB) was added, reactions were incubated at 37°C for 15 min, and 31 μl of 2x RNA loading dye was added.

Nucleic acids were separated by electrophoresis on 10% 7M urea slab gels (equilibrated and run in 1x TBE). Gels were incubated with SYBR Gold nucleic acid gel stain, bands were visualized with UV transillumination, and species ~80 to ~150 nt in length were recovered from the gel (procedure as above) and recovered in 20 μl of nuclease-free water.

#### cDNA amplification

cDNA derived from RNA products generated *in vitro* or *in vivo* were diluted with nuclease-free water to a concentration of ~10^9^ molecules/μl. 2μl of the diluted cDNA solution was used as a template for emulsion PCR reactions containing Illumina index primers using a Micellula DNA Emulsion and Purification Kit (EURx). The Illumina PCR forward primer and Illumina index primers from the TruSeq Small RNA Sample Prep Kits were used. The emulsion was broken, and DNA was purified according to the manufacturer’s recommendations. Amplicons were gel purified on 10% TBE slab gels (Invitrogen; equilibrated in 1x TBE), recovered by isopropanol precipitation and reconstituted in 13 μl of nuclease-free water.

#### High-throughput sequencing

Barcoded libraries were pooled and sequenced on an Illumina NextSeq platform in high-output mode using custom sequencing primer s1115.

#### Sample serial numbers

Samples KS112-KS114 are cDNA derived from RNA products generated in stationary-phase *E. coli* cells treated with (i) both PNK and Rpp (PNK + Rpp), (ii) Rpp only (Rpp), (iii) PNK only (PNK), or (iv) neither PNK nor Rpp (mock). Samples KS86-KS97 are cDNA derived from RNA products generated *in vitro* in the presence of no UpA (KS86-KS88), 40 μM UpA (KS89-KS91), 160 μM UpA (KS92-KS94), or 640 μM UpA (KS95-KS97) treated with (i) Rpp only (Rpp), (ii) PNK only (PNK), or (iii) neither PNK nor Rpp (mock).

#### Data analysis: separation of RNA 5’-end sequences by enzymatic treatment, promoter sequence, and promoter position

RNA 5’-end sequences were associated with an enzymatic treatment using the 4-nt barcode sequence acquired upon ligation of the 5’-adaptor (see above) as described in (16). RNA 5’-end sequences were associated with a p*lac*CONS promoter sequence using transcribed-region barcode assignments derived from the analysis of sample Vv945 described in (14). RNA 5’-end sequences that could be aligned to their template of origin with no mismatches were used for results presented in Figures 2, 3A, 4, S1, S5, and S6. RNA 5’-end sequences with mismatches at the first and/or second base of the 5’-end were also included for results shown in Figures 3B, S2–S3.

#### Data analysis: 5’-end distribution histograms (Figures 2, S1B, S8)

The number of 5’-end sequences emanating from each position 4 to 10 bp downstream of the −10 element of p*lac*CONS was determined for each of the ~4^10^ (~1,000,000) promoter sequences. These counts are represented using four vectors, 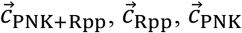, and 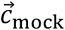, which represent the number of counts observed for 5’-ends at positions *i* = 4, 5, …, 10 for each enzymatic treatment. We initially estimated the number of 5’-ppp RNAs and 5’-OH RNAs in two different ways:

We initially computed the number of 5’-ppp RNAs and 5’-OH RNAs in two different ways:

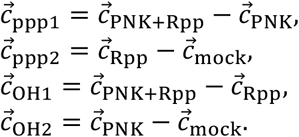

These four read count distributions were computed separately for each of the three replicates. To visualize these distributions, we normalized each counts vector by its sum across all positions, i.e.,

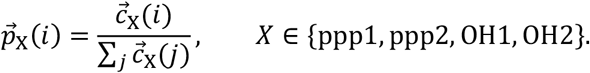

The 5’-OH distributions that resulted exhibited two obvious problems (Figure S8A, left): there was substantial variation across replicates, and four of the distributions exhibited probabilities well below zero at two positions (*i* = 8,9). We reasoned that these defects might be artefacts resulting from the enzymatic treatments not being 100% efficient, and that accounting for these inefficiencies might lead to more accurate 5’-OH distributions. Let *ϵ*_PNK+Rpp_, *ϵ*_PNK_, *ϵ*_Rpp_, and *ϵ*_mock_ denote the efficiencies of the four enzymatic treatments. Then the number of true underlying counts in the four samples becomes,

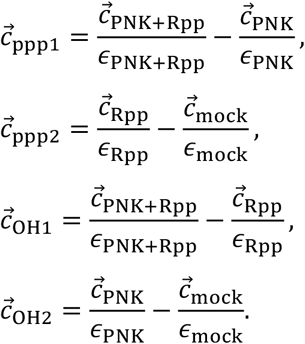

One can solve for the unknown efficiencies by setting 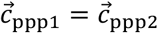, or equivalently 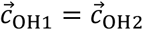, both of which give

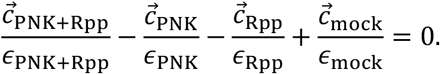

This provides a system of 7 equations with 4 unknowns, and setting *ϵ*_mock_ = 1 allowed us to solve for the other three other (now relative) efficiencies. We did this by minimizing the objective function

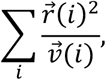

where

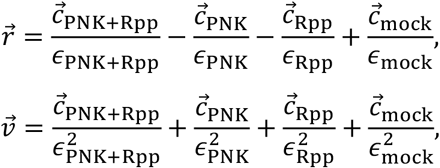

are vectors that respectively represent the residuals and Poisson-estimated variances (Figure S8B). Using these efficiencies, we computed the corrected read count distributions (Figure S8A, right). The resulting 5’-OH distributions were much more reproducible across replicates. Moreover, a negative probability was estimated only for position 9 for 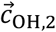 of replicate 2, and even this was much closer to zero than in the uncorrected profiles (Figure S8A, right). We therefore chose to compute 5’-ppp and 5’-OH read counts using the averages,

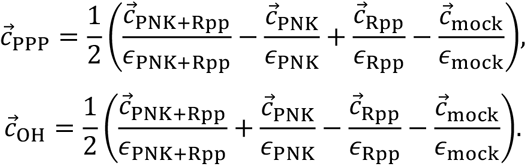

These averaged distributions were used in the *in vivo* MASTER analysis shown in Figures 2 and S1A, as were analogous formulas for analyzing primer usage and for generating the sequence logos shown in Figures 3–4 and S5.

#### Data analysis: modeling 5’-OH distributions as a mixture of shifted 5’-ppp distributions (Figure S1A)

We modeled the 5’-end distributions of 5’-OH RNAs as a mixture of 5’-ppp RNAs, each shifted between −4 nt and +4 nt. The mixture coefficients were inferred using least-squares regression under positivity and normalization constraints. The 5’-ppp distributions shifted one nucleotide upstream account for 78.7% ± 4.8% of the mixture model, no shift in the 5’-ppp distributions accounts for 20.0% ± 4.8% of the mixture model, and all other shifts account for 0.39% ± 0.06% of the model (uncertainties represent the SD across replicates).

To carry out the mixture modeling of 5’-end distributions, we defined a 7×9 matrix *A*, whose entries *A*_*ij*_ represent the fraction of reads with 5’-ends at positions 4-10 (corresponding to *i* = 1, 2, …, 7 within the 5’-ppp distribution shifted by −4, −3, …, +4 nt (corresponding to *j* = 1, 2, …, 9). Note that these shifted distributions were normalized so that ∑_*i*_ *A*_*ij*_ = 1 for every column *j*. We further defined a 7×1 vector 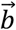 representing the 5’-OH distribution, normalized so that ∑_*i*_ *b*_*i*_ = 1. We then inferred a 9×1 vector of mixture coefficients 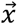 by solving the constrained least squares problem

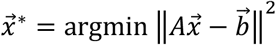

under the constraints that all *x*_*j*_ ≥ 0 and that ∑_*j*_ *x*_*j*_ = 1. The resulting mixture distribution 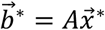 is shown in Figure S1A (left panel), alongside the 5’-OH distribution 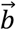. The corresponding mixture coefficients are shown in Figure S1A (right panel). The residual deviation was computed as

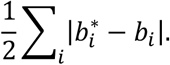

#### Data analysis: in vivo sequence logos (Figures 4, S5)

Sequence logos illustrate the sequence-dependent log_2_ likelihood of primer-dependent initiation for primer binding sites 6-7, 7-8, and 8-9. All logos were created using Logomaker (35). The specific quantities illustrated in the logos were computed as follows.

To generate the logos shown in Figures 4 (left) and S5, we first estimated the values of 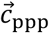 and 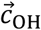 originating from promoters for which each base *b* ∈ {*A*, *C*, *G*, *T*} occurs at each position *l* ∈ {1,2, …, 10} within the 10 bp randomized region (i.e., positions 1-10 bp downstream of the −10 element). These read count estimates were computed as

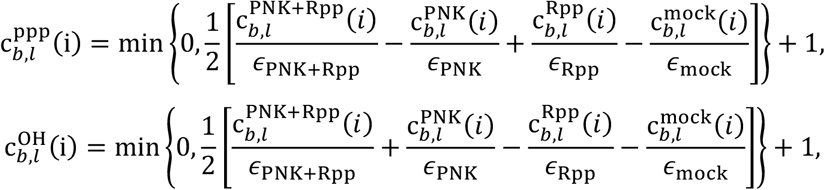

where *i* ∈ {4, 5, …, 10} indicates the location of the 5’ end of the tallied transcripts, and 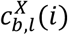 is the number of transcripts observed to originate at position *i* from promoters with base *b* at position *l* in the sample corresponding to enzymatic treatment *X*. The minimum is to prevent counts from becoming negative, and the “+ 1” is a pseudo-count used to regularize the computation of log ratios. We then computed the total number of such reads summed over 5’-end positions *i*:

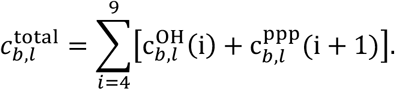

A logo for each 5’-end position *i* was then plotted, where the height of each character *b* at each position *l* was given by the centered log_2_ ratio

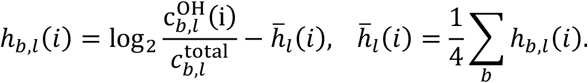

Logos were computed separately for each of the three biological replicates. Logos shown in Figure 4 (left panel) were created using reads from all promoter sequences. Logos shown in Figures S5 and S6 were created using reads from promoter sequences with the specified primer binding site at positions *i* and *i* + 1.

#### Data analysis: in vitro sequence logos (Figures 4, S6)

To generate the sequence logos shown in Figures 4 (right) and S6, we estimated the values of 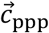 and 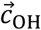 as

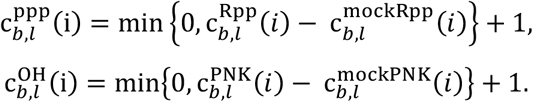

These estimated counts were used to generate logos for promoter sequences with a UpA binding site at positions *i* and *i* + 1 using the procedure described above.

#### Data analysis: dinucleotide usage (Figure 3A)

The percent usage of each dinucleotide primer *x* ∈ {ApA, ApC, …, UpU} was computed as

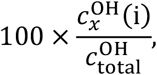

where

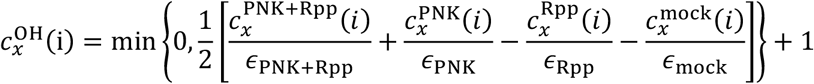

is the regularized efficiency-corrected count of 5’-OH RNAs originating from promoters with a binding site for primer *x* at positions *i* and *i* + 1, and

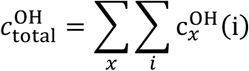

is the sum of such counts over positions and primers.

#### Data analysis: primer binding site complementarity (Figures 3B, S2, S3)

To compute the binding site complementarity results, we first computed the efficiency-corrected regularized read counts

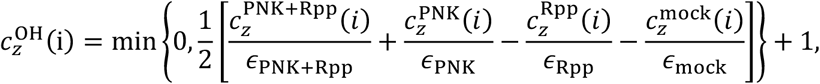

which are conditioned on the indicator variable

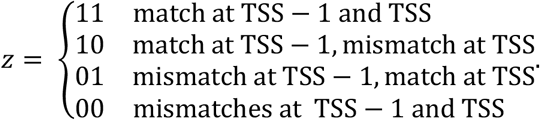

The corresponding total counts were computed as

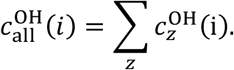

The position-dependent “% 5’-OH RNA” values plotted in Figures S2 and S3 are given by

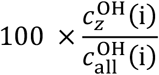

for appropriate choice of *z*. The corresponding percentages aggregated across positions, shown in Figures 3B and S3, are given by

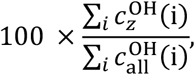

where the sums are over positions 6, 7, and 8.

### Analysis of primer-dependent initiation from chromosomally-encoded *E. coli* promoters (Figure 6A, Table S1)

#### cDNA library construction and sequencing

Cell growth, RNA isolation, enzymatic treatments, and 5’-adaptor ligations were performed as described above. 25 μl of 5’-adaptor-ligated RNAs were mixed with 5 μl of 18.5 μM (1.86 μM final) of an oligo pool consisting of a mixture of 93 gel purified oligodeoxyribonucleotides each having 5’-end sequence identical to the “RT primer” contained in Illumina TruSeq Small RNA Sample Prep Kits and a 3’-end sequence complementary to a chromosomally-encoded *E. coli* promoter that uses UpA as primer (Table S3). The mixture was incubated at 65°C for 5 min, kept at 4°C, combined with 20 μl of a solution containing 10 μl of 5x First-Strand buffer (Invitrogen), 2.5 μl of 10 mM dNTP mix (NEB), 2.5 μl of 100 mM DTT (Invitrogen), 2.5 μl of 40 U/μl RNaseOUT, and 2.5 μl of 100 U/μl SuperScript III Reverse Transcriptase (Invitrogen), for a final reaction volume of 50 μl. Reactions were incubated at 25°C for 5 min, 55°C for 60 min, 70°C for 15 min, then cooled to 25°C. Next, 5.4 μl of 1M NaOH was added, reactions were incubated at 95°C for 5 min, cooled to 10°C, 4.5 μl of 1.2M HCl was added, followed by 60 μl of 2x RNA loading dye. Nucleic acids were separated by electrophoresis on 10% 7M urea slab gels (equilibrated and run in 1x TBE). Gels were incubated with SYBR Gold nucleic acid gel stain, bands were visualized with UV transillumination, and species ~80 to ~150 nt in length were recovered from the gel (procedure as above) and recovered in 20 μl of nuclease-free water.

cDNA amplification and high-throughput sequencing was performed as described above. Serial numbers for these samples are KS118-KS120.

#### Data analysis: chromosomal promoter sequence logo (Figure 6A)

Sequencing reads were associated with one of the four reaction conditions based on the identity of the 4-nt barcode sequence. RNA 5’-end sequences that could be aligned to the chromosomally-encoded promoter from which they were expressed with no mismatches were used for results presented in Figure 6A. The number of 5’-end sequences emanating from each position up to four bases upstream and downstream of the UpA binding site (TSS-5 to TSS+4, where UpA binds positions TSS-1, TSS) was determined for each enzymatic treatment. To represent these data as a sequence logo, we first estimated the fraction of transcripts that had 5′-OH ends at TSS-1. For each promoter sequence *s*, this was computed using

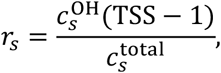

where

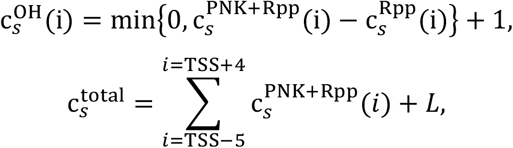

and where 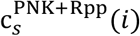 and 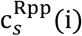 denote the number of read initiating from position *i* (which ranges over *L*=10 positions, from TSS-5 to TSS+4) observed for promoter *s* in the PNK+Rpp and Rpp treatments, respectively. These *r*_*s*_ values were then averaged across three replicates, and a sequence logo reflecting the average log_2_ value of these ratios was rendered using mean-centered character heights given by

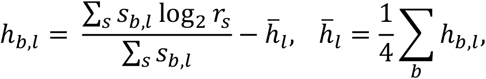

where *s*_*b,l*_ takes the value 1 if base *b* occurs at position *l* in sequence *s* and is 0 otherwise.

### Single-template *in vitro* transcription assays (Figures 5, S4)

10 nM of linear template was mixed with 50 nM RNAP holoenzyme in transcription buffer and incubated at 37°C for 15 min to form open complexes. 1000 μM ATP and increasing concentrations of UpA (0, 10, 40, 160, and 640 μM) were added along with 10 μM of non-radiolabeled UTP plus 6 mCi of [α^32^P]-UTP (PerkinElmer). Upon addition of nucleotides, reactions were incubated at 37°C for 10 min to allow for product formation. Reactions were stopped by addition of an equal volume of gel loading buffer (95% formamide; 25 mM EDTA; 0.025% SDS, 0.025% xylene cyanol; 0.025% bromophenol blue).

Samples were run on 20% TBE-Urea polyacrylamide gels. Bands were quantified using ImageQuant software. Observed values of UpApU / (pppApU + UpApU) were plotted vs. [UpA] / [ATP] on semi-log plot (Sigmaplot). Non-linear regression was used to fit the data to the equation: y = (ax) / (b+x); where y is UpApU / (pppApU + UpApU), x is [UpA] / [ATP], and a and b are regression parameters. The resulting fit yields the value of x for which y = 0.5. The relative efficiency (k_cat_/K_M_)_UpA_ / (k_cat_/K_M_)_ATP_ is equal to 1/x.

### Analysis of primer-dependent initiation from the *E. coli bhsA* promoter (Figure 6B)

#### Culture growth and cell harvesting

Plasmids pBEN516, pKS494 or pKS497 were introduced into *E. coli* MG1655 cells. Plasmid-containing cells were grown in 25 ml of LB containing chloramphenicol (25 μg/ml) in a 125 ml DeLong flask (Bellco Glass) at 37°C and harvested 5, 9, 14, or 21 h after cells had entered stationary phase (OD600 of ~3.3, ~3.1, ~2.9, and ~2.6, respectively). Cell suspensions were removed to 2 ml microcentrifuge tubes (Axygen), cells were collected by centrifugation (15,000 rpm; 30 s; 4°C), and cell pellets were stored at −80°C.

#### RNA isolation

Cells were resuspended in 0.6 ml of TRI-Reagent, incubated at 70°C for 10 min, and the cell lysate was centrifuged to remove insoluble material (10 min; 21,000 x *g*; 4°C). The supernatant was transferred to a fresh tube, ethanol was added to a final concentration of 60.5%, and the mixture was applied to a Direct-zol spin column. DNase I treatment was performed on-column according to the manufacturer’s recommendations. RNA was eluted from the column using nuclease-free water that had been heated to 70°C (3 × 30 μl elutions; total volume of eluate = 90 μl). RNA was treated with 2 U TURBO DNase at 37°C for 1 h to remove residual DNA. Samples were extracted with acid phenol:chloroform, RNA was recovered by ethanol precipitation and resuspended in RNase-free water.

#### Primer extension analysis

Assays were performed essentially as described in (10). 10 μg of RNA was combined with 3 μM of primer k711 (5’-radiolabeled using PNK and [γ^32^P]-ATP). The RNA-primer mixture was heated to 95°C for 10 min, slowly cooled to 40°C (0.1°C/s), incubated at 40°C for 10 min, and cooled to 4°C using a thermal cycler (Biorad). Next, 10 U of AMV reverse transcriptase (NEB) was added, reactions were incubated at 55°C for 60 min, heated to 90°C for 10 min, cooled to 4°C for 30 min, and mixed with 10 μl of 2x RNA loading buffer (95% formamide; 0.5 mM EDTA, pH 8; 0.025% SDS; 0.0025% bromophenol blue; and 0.0025% xylene cyanol). Nucleic acids were separated by electrophoresis on 8%, 7M urea slab gels (equilibrated and run in 1x TBE) and radiolabeled products were visualized by storage phosphor imaging. Band assignments were made by comparison to a DNA sequence ladder prepared using primer k711 and pBEN516 as template (Affymetrix Sequenase DNA sequencing kit, version 2).

### Structure determination (Figures 7, S7, Table S2)

The nucleic-acid scaffold for assembly of *Thermus thermophilus* RPo was prepared from synthetic oligonucleotides (Sangon Biotech) by an annealing procedure (95°C, 5 min followed by 2°C-step cooling to 25°C) in 5 mM Tris-HCl (pH 8.0), 200 mM NaCl, and 10 mM MgCl_2_ (nontemplate strand for all structures: 5’-TATAATGGGAGCTGTCACGGATGCAGG-3’; template strand for RPo[A_TSS-2_A_TSS-1_T_TSS_]-UpA-CMPcPP: 5’-CCTGCATCCGTGAGTAAAG-3’; template strand for RPo[A_TSS-2_C_TSS-1_C_TSS_]-GpG-CMPcPP: 5’-CCTGCATCCGTGAGCCAAG-3’; template strand for RPo[T_TSS-2_C_TSS-1_C_TSS_]-GpG-CMPcPP: 5’-CCTGCATCCGTGAGCCTAG-3’).

For each structure, *T. thermophilus* RPo was reconstituted by mixing *T. thermophilus* RNAP holoenzyme purified as described in (18) and the nucleic-acid scaffold at a 1:1.2 molar ratio and incubating for 1h at 22°C. Crystals of *T. thermophilus* RPo were obtained and handled essentially as in (18). The primer (UpA or GpG) and CMPcPP were subsequently soaked into RPo crystals by addition of 0.2 μl 100 mM primer (UpA or GpG) and 0.2 μl 50 mM CMPcPP in RNase-free water to the crystallization drops (2 μl) and incubation for 30 min at 22ºC. Crystals were transferred in a stepwise fashion to reservoir solution (0.2 M KCl; 0.05 M MgCl_2_; 0.1 M Tris-HCl, pH 7.9; 9% PEG 4000) containing 0.5%, 1%, 5%, 10%, and 17.5% (v/v) (2R, 3R)-(−)-2,3-butanediol, and cooled in liquid nitrogen.

Diffraction data were collected at Shanghai Synchrotron Radiation Facility (SSRF) beamlines 17U and processed using HKL2000 (36). The structures were solved by molecular replacement with Phaser MR in Phenix using one molecule of RNAP holoenzyme from the structure of *T. thermophilus* RPo (PDB:4G7H) as the search model (18, 37). Early-stage rigid-body refinement of the RNAP molecule revealed good electron density signals for the primer (UpA or GpG) and CMPcPP. Cycles of iterative model building with Coot and refinement with Phenix were performed (38, 39). The models of the primer (UpA or GpG) and CMPcPP were built into the map at later stage of refinement.

The final model of RPo[A_TSS-2_A_TSS-1_T_TSS_]-UpA-CMPcPP was refined to R_work_ and R_free_ of 0.205 and 0.245, respectively. The final model of RPo[A_TSS-2_C_TSS-1_C_TSS_]-GpG-CMPcPP was refined to R_work_ and R_free_ of 0.187 and 0.252, respectively. The final model of RPo[T_TSS-2_C_TSS-1_C_TSS_]-GpG was refined to R_work_ and R_free_ of 0.194 and 0.246, respectively.

### Quantification and statistical analysis

The number of replicates and statistical test procedures are in the figure legends.

### Data and software availability

Sequencing reads have been deposited in the NIH/NCBI Sequence Read Archive under the study accession number PRJNA718578. Logos were generated using Logomaker (35) via custom Python scripts. Source code and documentation are provided at http://www.github.com/jbkinney/20_nickels. Structures of RPo[A_TSS-2_A_TSS-1_T_TSS_]-UpA-CMPcPP, RPo[A_TSS-2_C_TSS-1_C_TSS_]-GpG-CMPcPP, and RPo[T_TSS-2_C_TSS-1_C_TSS_]-GpG have been deposited in the Protein Data Bank (PBD) under identification codes 7EH0, 7EH1, and 7EH2, respectively.

## Acknowledgements

Work was supported by National Natural Science Foundation of China grant 31822001 (YZ) and National Institutes of Health grants GM124976 (PS), GM133777 (JBK), GM041376 (RHE), and GM118059 (BEN).

---

**Figure S1.**
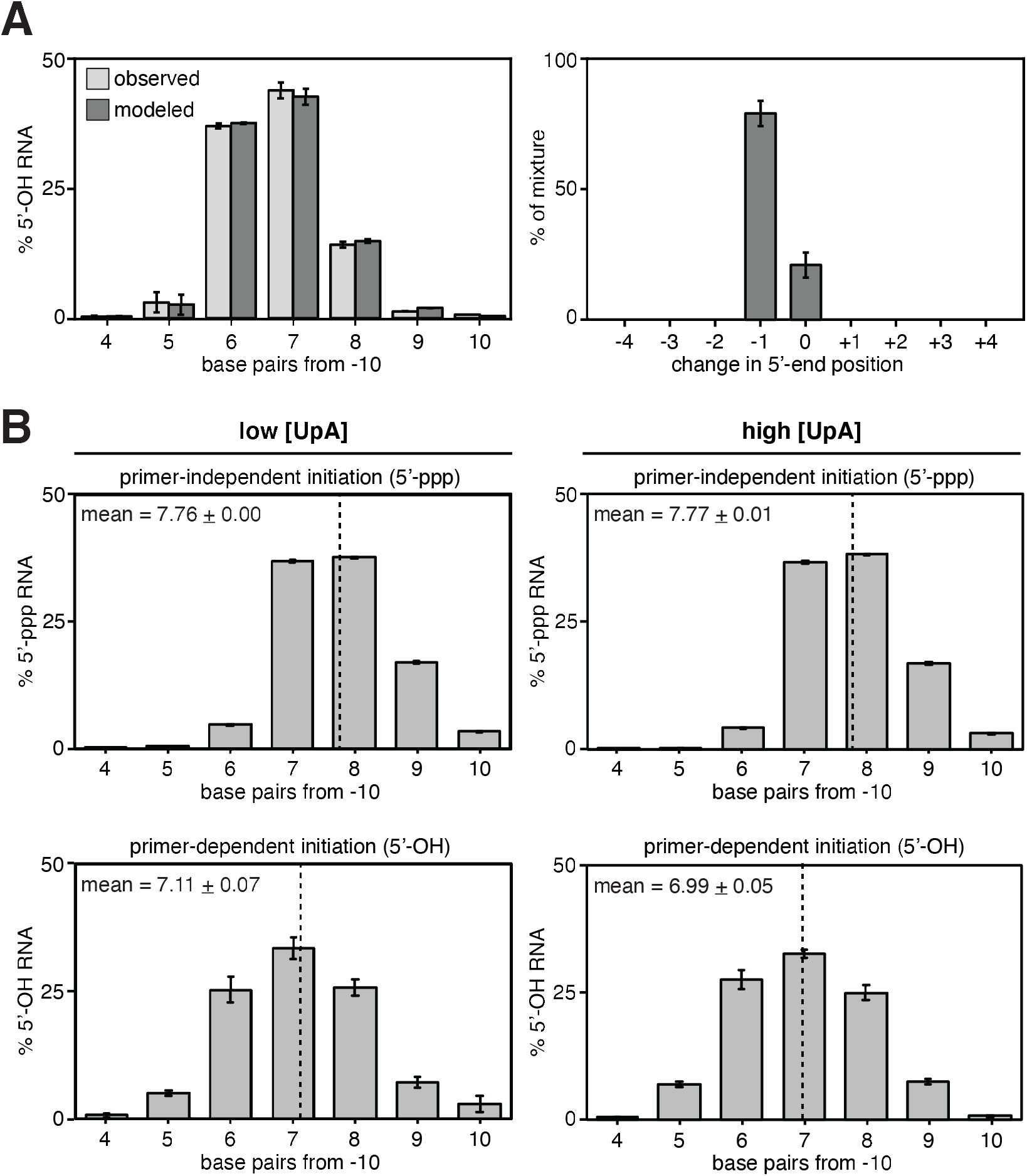
Distributions of 5′-end sequences for RNAs generated in primer-independent initiation and primer-dependent initiation. **A.** Computational modeling of *in vivo* distributions of 5′-OH RNAs from observed *in vivo* distributions of 5′-ppp RNAs (see Figure 2B, top). Left: histograms of observed *in vivo* distributions of 5′-OH RNAs (light gray; see also Figure 2B, bottom) and modeled distributions of 5′-OH RNAs (dark gray). The modeled distributions of 5′-OH RNAs was generated by modeling distributions of 5′-OH RNAs as a mixture of 5′-ppp RNAs with positions changed by up to 4 bp upstream (−1 to −4) or downstream (+1 to +4). Right: Mixture coefficient histogram. Coefficients were inferred using least-squares regression under positivity and normalization constraints. **B.** RNA 5′-end distribution histograms (mean ± SD, N = 3) for RNAs generated by primer-independent initiation or primer-dependent initiation *in vitro* (top and bottom, repectively). low UpA, [UpA] / [NTPs] = 0.04; high UpA, [UpA] / [NTPs] = 0.64. Dashed line indicates the mean 5′-end position (mean ± SD, N = 3).

**Figure S2.**
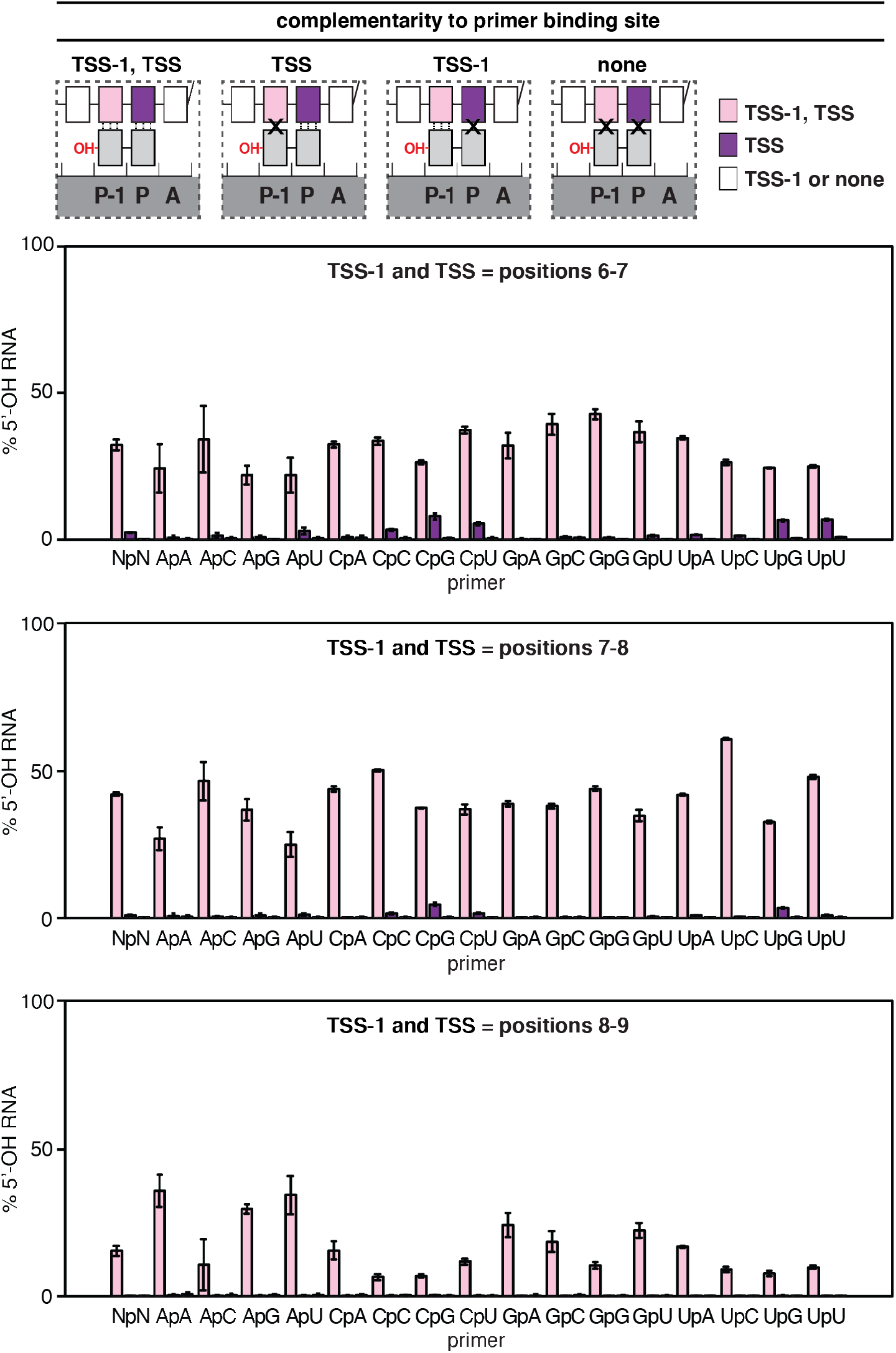
Promoter-sequence dependence of primer-dependent initiation in stationary-phase *E. coli* cells: primer binding site. Top: primer-dependent initiation involving template-strand complementarity to both 5′ and 3′ nucleotides of primer (TSS-1, TSS), template-strand complementarity to only 3′ nucleotide of primer (TSS), template-strand complementarity to only 5′ nucleotide of primer (TSS-1), or no template-strand complementarity to primer (none). Three vertical lines, complementarity; X, non-complementarity. Other symbols and colors as in Figure 1. Bottom: percentage of primer-dependent initiation involving complementarity to both 5′ and 3′ nucleotides of primer (TSS-1, TSS; pink), complementarity to only 3′ nucleotide of primer (TSS; purple), or template-strand complementarity to only 5′ nucleotide of primer or no template-strand complementarity to primer (TSS-1 or none; white) in stationary-phase *E. coli* cells for primer binding sites located 6-7, 7-8, or 8-9 base pairs downstream of the promoter −10 element (mean ± SD, N = 3).

**Figure S3.**
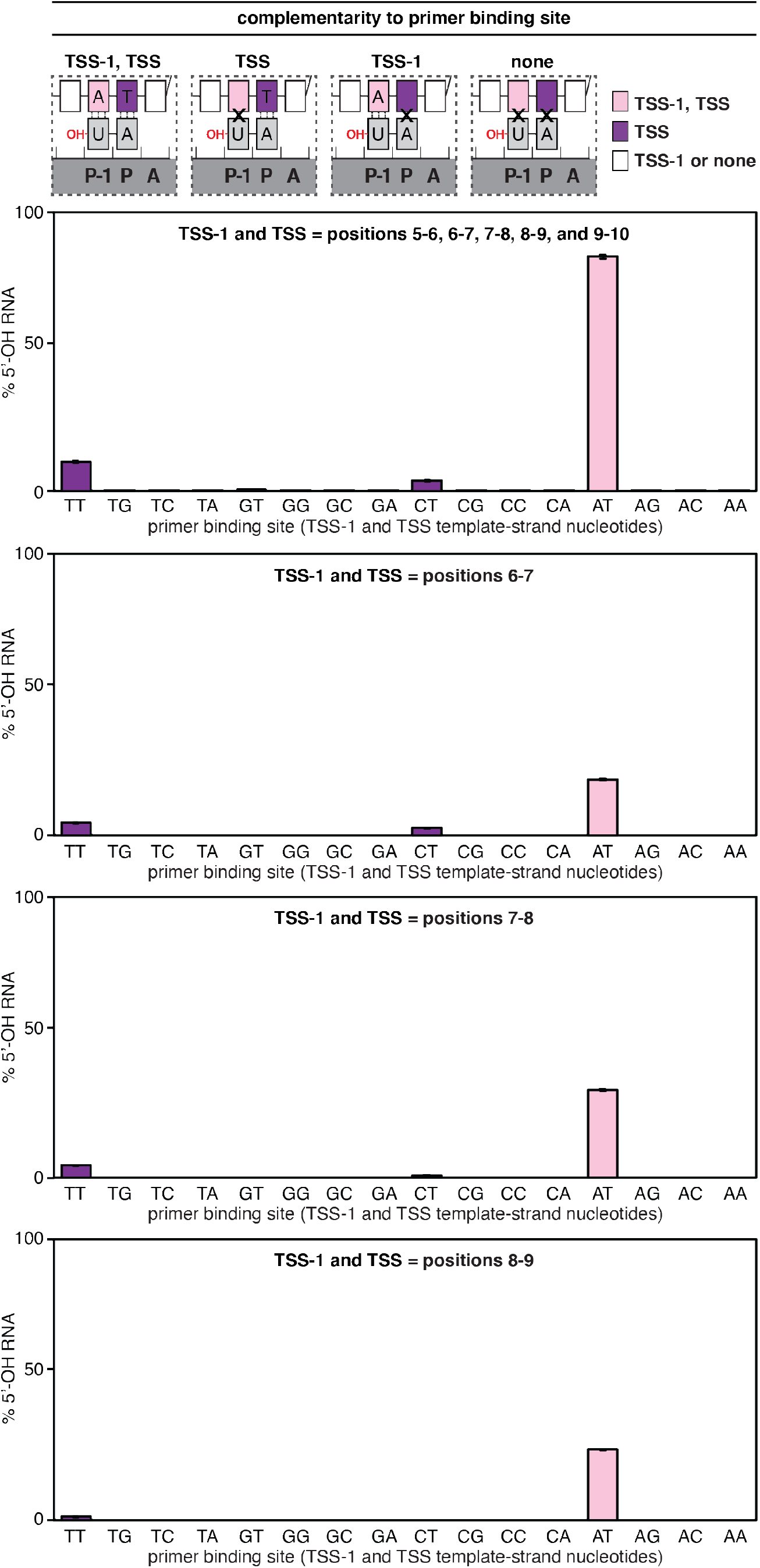
Promoter-sequence dependence of primer-dependent initiation *in vitro*: primer binding site. Top: primer-dependent initiation involving template-strand complementarity to both 5′ and 3′ nucleotides of UpA (TSS-1, TSS), template-strand complementarity to only 3′ nucleotide of UpA (TSS), template-strand complementarity to only 5′ nucleotide of UpA (TSS-1), or no template-strand complementarity to UpA (none). Three vertical lines, complementarity; X, non-complementarity. Other symbols and colors as in Figure 1. Bottom: percentage of primer-dependent initiation involving complementarity to both 5′ and 3′ nucleotides of UpA (TSS-1, TSS; pink), complementarity to only 3′ nucleotide of UpA (TSS; purple), or template-strand complementarity to only 5′ nucleotide of UpA or no template-strand complementarity to UpA (TSS-1 or none; white) for primer binding sites located 6-7, 7-8, or 8-9 base pairs downstream of the promoter −10 element (mean ± SD, N = 3).

**Figure S4.**
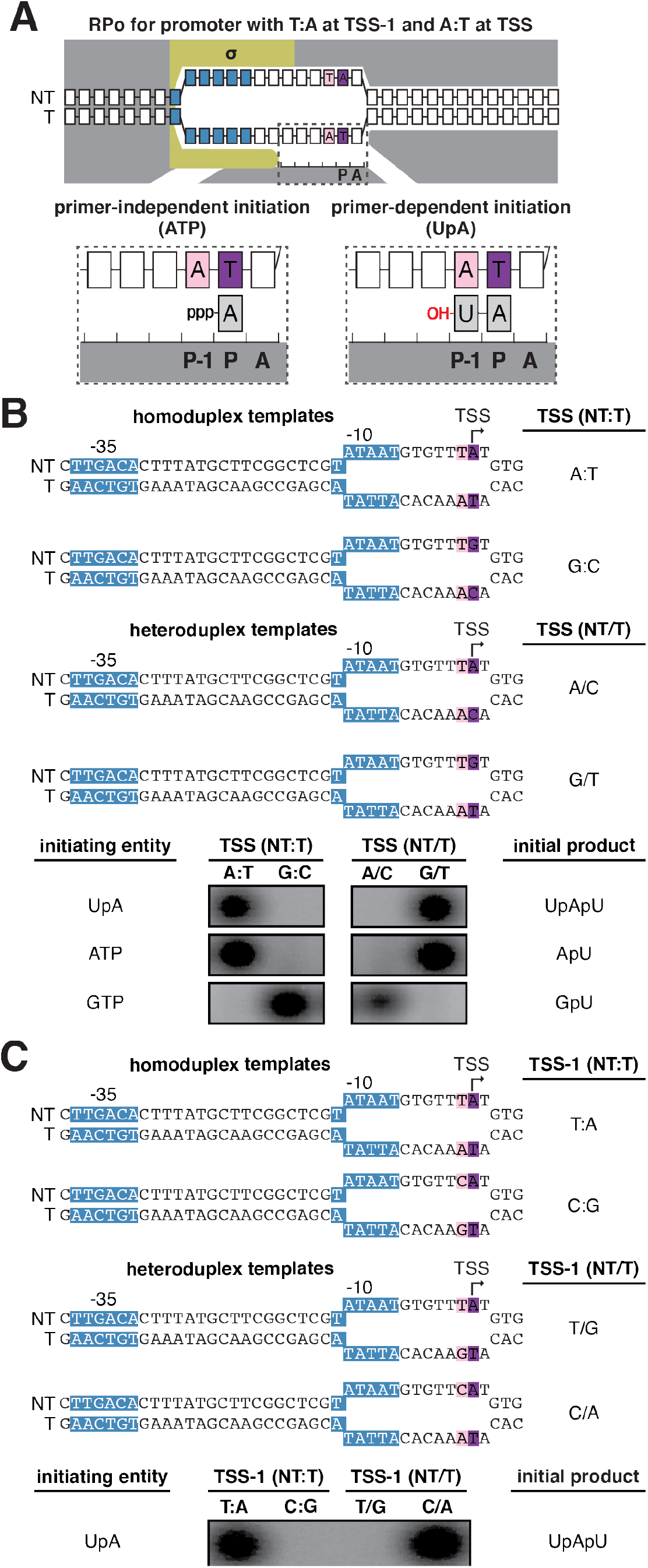
Promoter-sequence dependence of primer-dependent initiation *in vitro*: template strand carries sequence information at positions TSS and TSS-1. **A.** Binding of ATP or UpA to template-strand nucleotides in primer-independent initiation (left) and primer-dependent initiation (right). Unwound transcription bubble in RPo indicated by raised and lowered nucleotides. Other symbols and colors as in Figure 1. **B-C.** Strand specificity at positions TSS and TSS-1. Top: homoduplex DNA templates. Middle: heteroduplex DNA templates containing mismatches at positions TSS (panel A) or TSS-1 (panel B). Unwound transcription bubble in RPo indicated by raised and lowered nucleotides. Bottom: radiolabeled initial RNA products generated using the indicated template in reactions containing UpA, ATP or GTP (panel B) or UpA (panel C).

**Figure S5.**
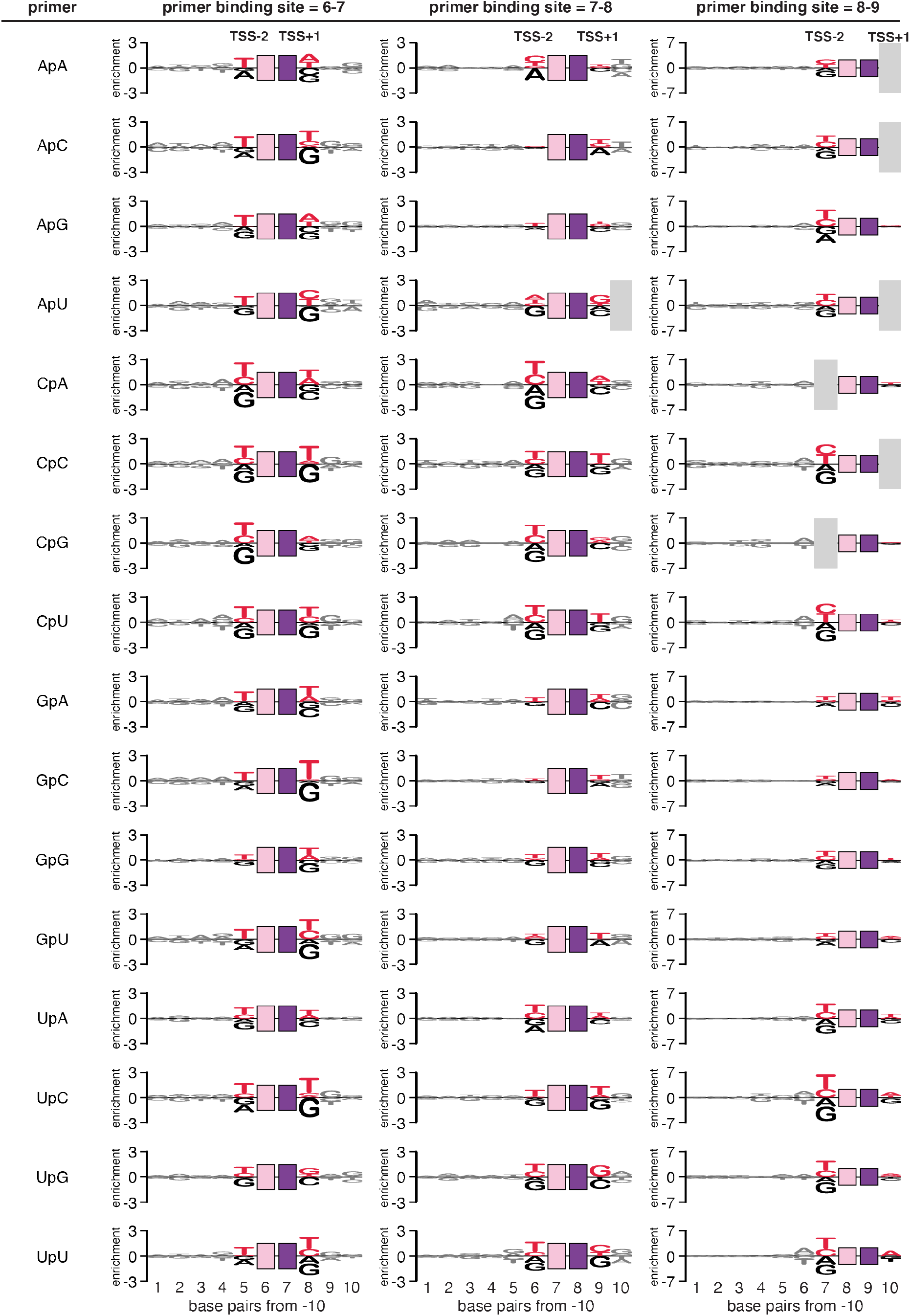
Promoter-sequence dependence of primer-dependent initiation *in vivo*: sequences flanking the primer binding site. Sequence logo (35) for primer-dependent initiation in stationary-phase *E. coli* cells with each of the 16 dinucleotides at TSS positions 7, 8, and 9 (corresponding to primer binding sites 6-7, 7-8, and 8-9, respectively). The height of each base “X” at each position “Y” represents the log_2_ average of the % 5′-OH RNAs computed across sequences containing nontemplate-strand X at position Y. Red, consensus nucleotides; black, non-consensus nucleotides. Gray box indicates positions where enrichment values could not be computed.

**Figure S6.**
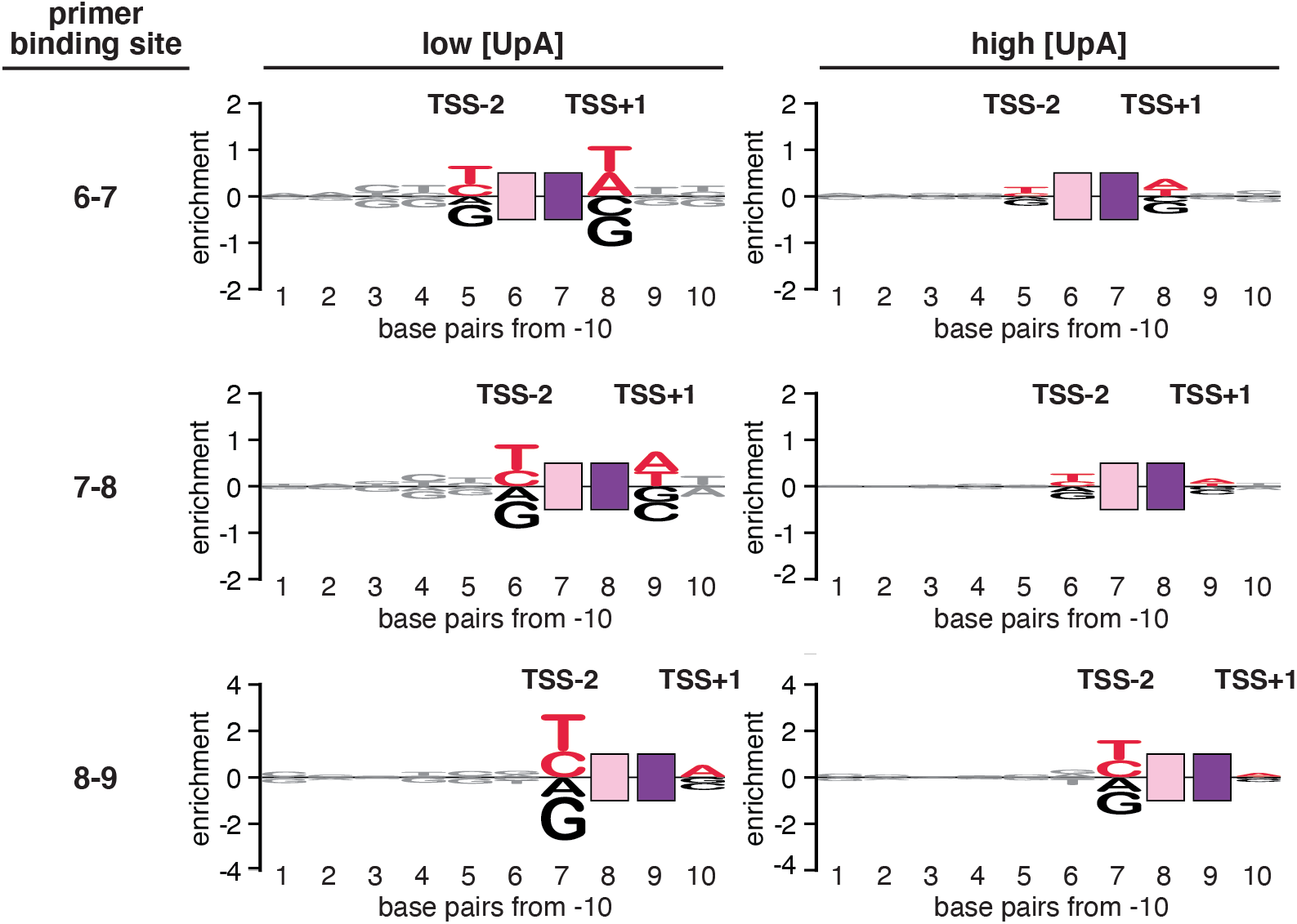
Promoter-sequence dependence of primer-dependent initiation *in vitro*: sequences flanking the primer binding site. Sequence logo (35) for primer-dependent initiation *in vitro* with UpA at TSS positions 7, 8, and 9 (corresponding to primer binding sites 6-7, 7-8, and 8-9, respectively). The height of each base “X” at each position “Y” represents the log_2_ average of the % 5′-OH RNAs computed across sequences containing nontemplate-strand X at position Y. Low UpA, [UpA] / [NTPs] = 0.04; high UpA, [UpA] / [NTPs] = 0.64. Red, consensus nucleotides; black, non-consensus nucleotides.

**Figure S7.**
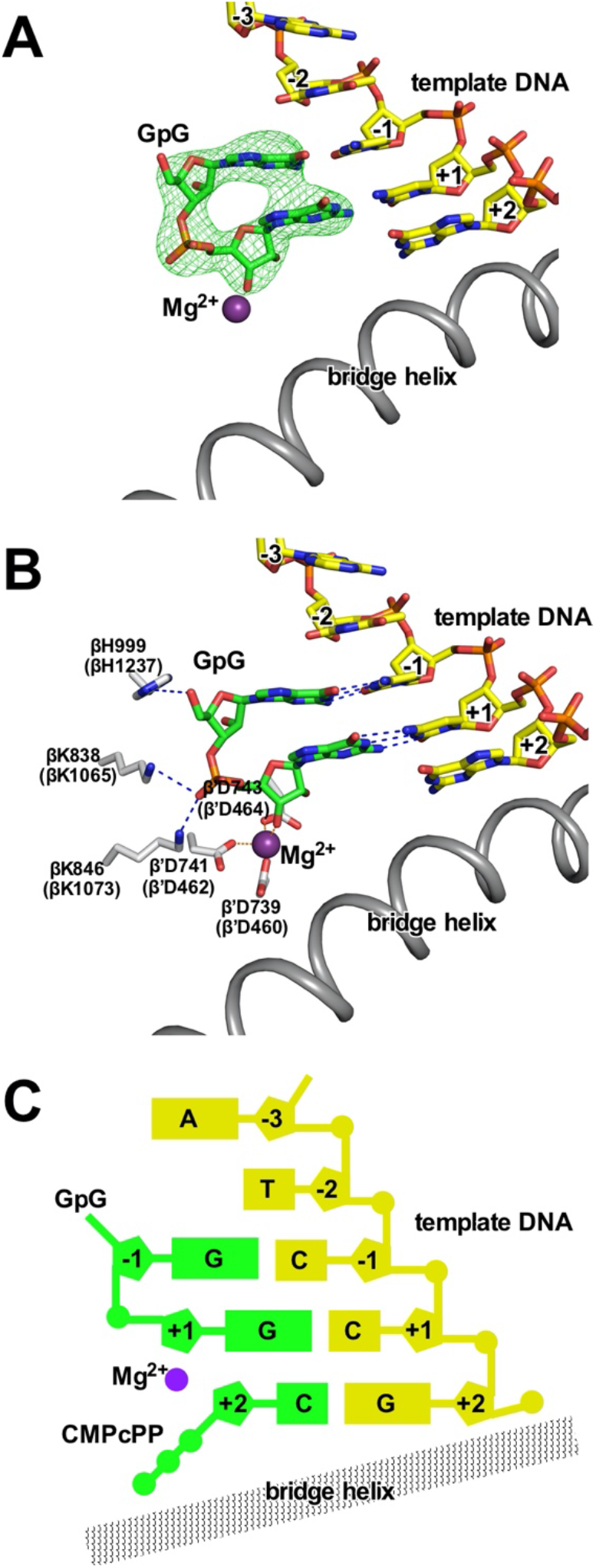
Structural basis of promoter-sequence dependence of primer-dependent initiation at position TSS-2. Crystal structure of *T. thermophilus* RPo[T_TSS-2_C_TSS-1_C_TSS_]-GpG-CMPcPP **A.** Experimental electron density (contoured at 2.5σ; green mesh) and atomic model for DNA template strand (yellow, red, blue, and orange for C, O, N, and P atoms), dinucleotide primer (green, red, blue, and orange for C, O, N, and P atoms), RNAP active-center catalytic Mg^2+^(I) (violet sphere), and RNAP bridge helix (gray ribbon). **B.** Contacts of RNAP residues (gray, red, and blue for C, O, and N atoms) with primer and RNAP active-center catalytic Mg^2+^(I). RNAP residues are numbered both as in *T. thermophilus* RNAP and as in *E. coli* RNAP (in parentheses). **C.** Schematic summary of structures. Template-strand DNA (yellow); primer (green); RNAP bridge helix (gray); RNAP active-center catalytic Mg^2+^(I) (violet). Note that, in contrast to structures with template-strand purine at position TSS-2 (Figure 7), in this structure with template-strand pyrimidine at position TSS-2, no density for CMPcPP is observed.

**Figure S8.**
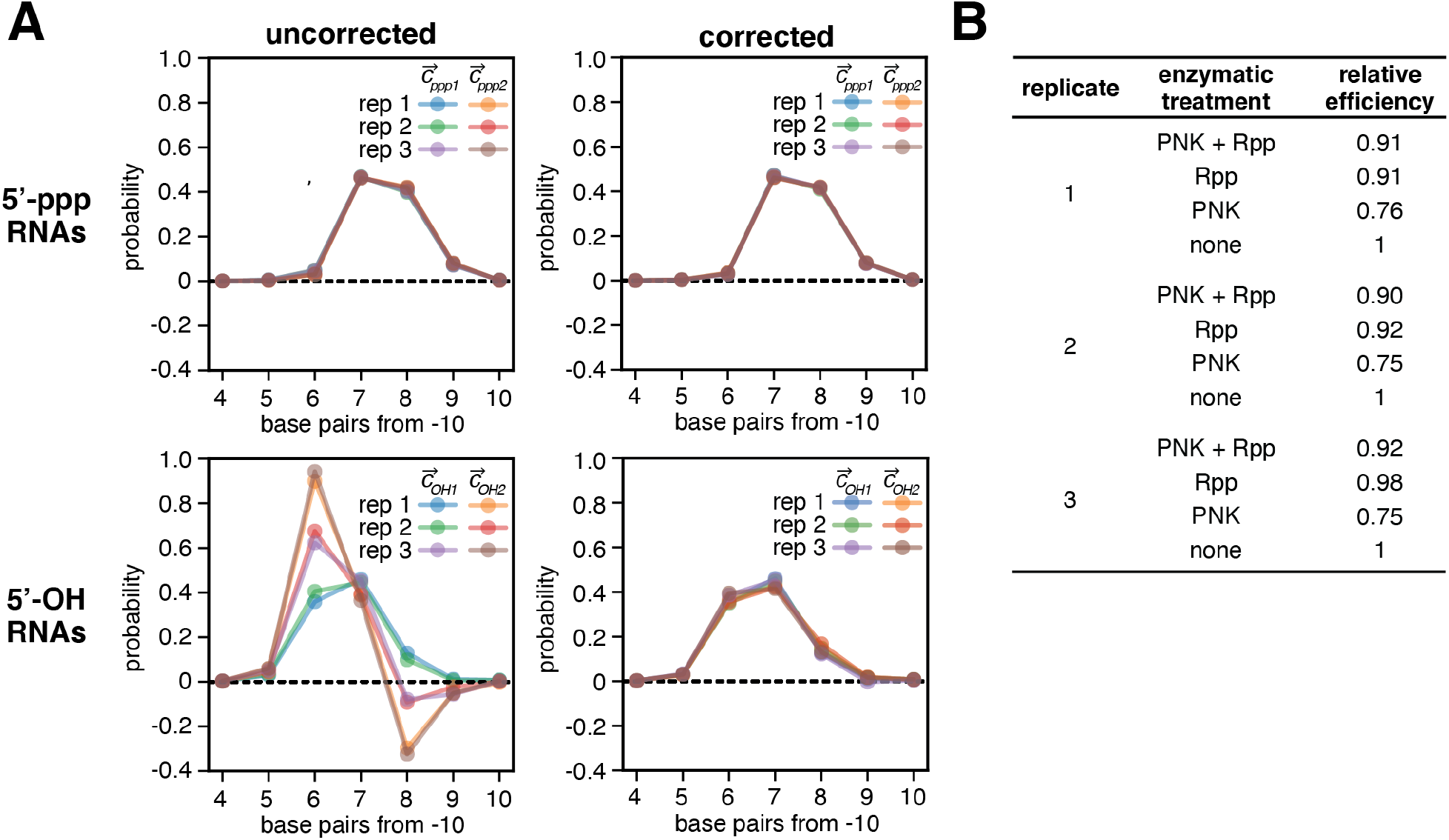
Primer-dependent initiation *in vivo*: MASTER data analysis. **A.** 5’-ppp distributions (top panels) and 5’-OH distributions (bottom panels) calculated using uncorrected read counts (left) or using read counts computed using correction factors that account for inefficiencies of enzymatic processing (corrected, right). **B.** Relative enzymatic processing efficiencies computed for each replicate.

**Table S1.**
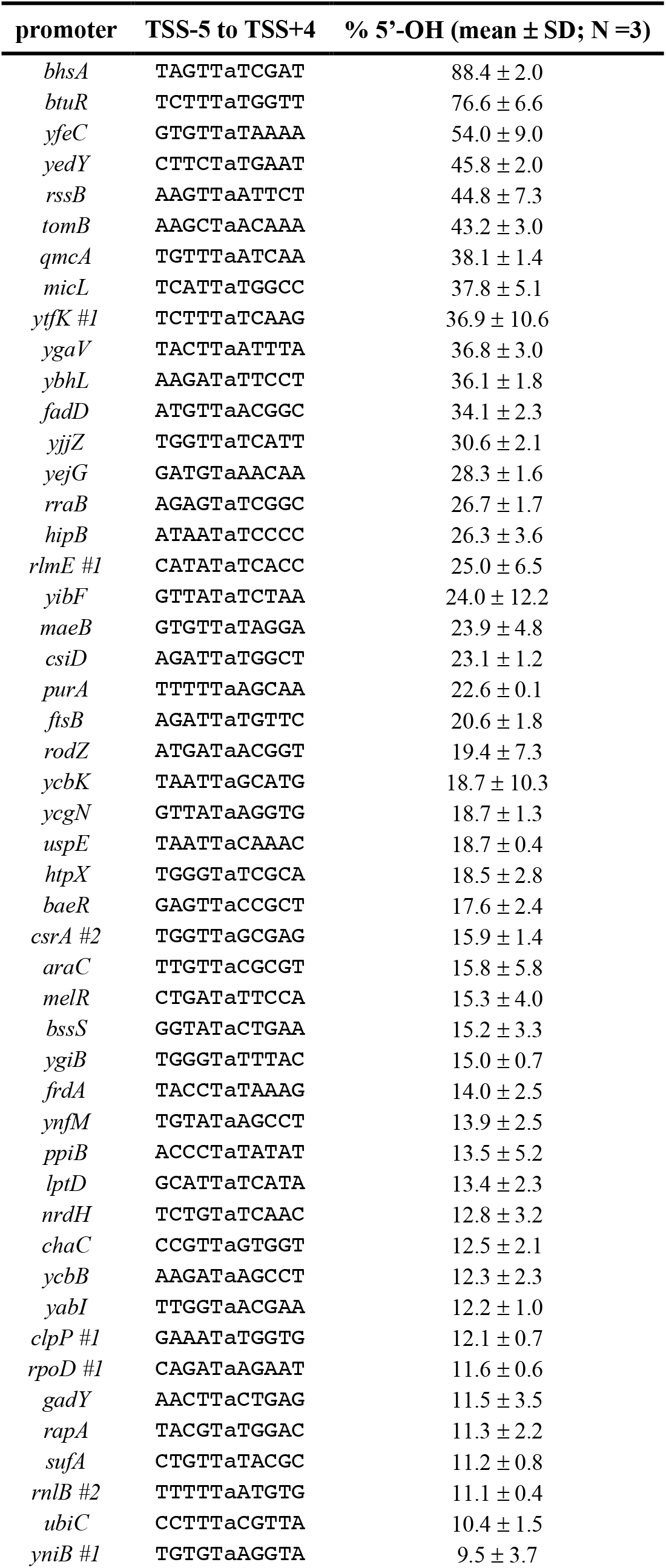
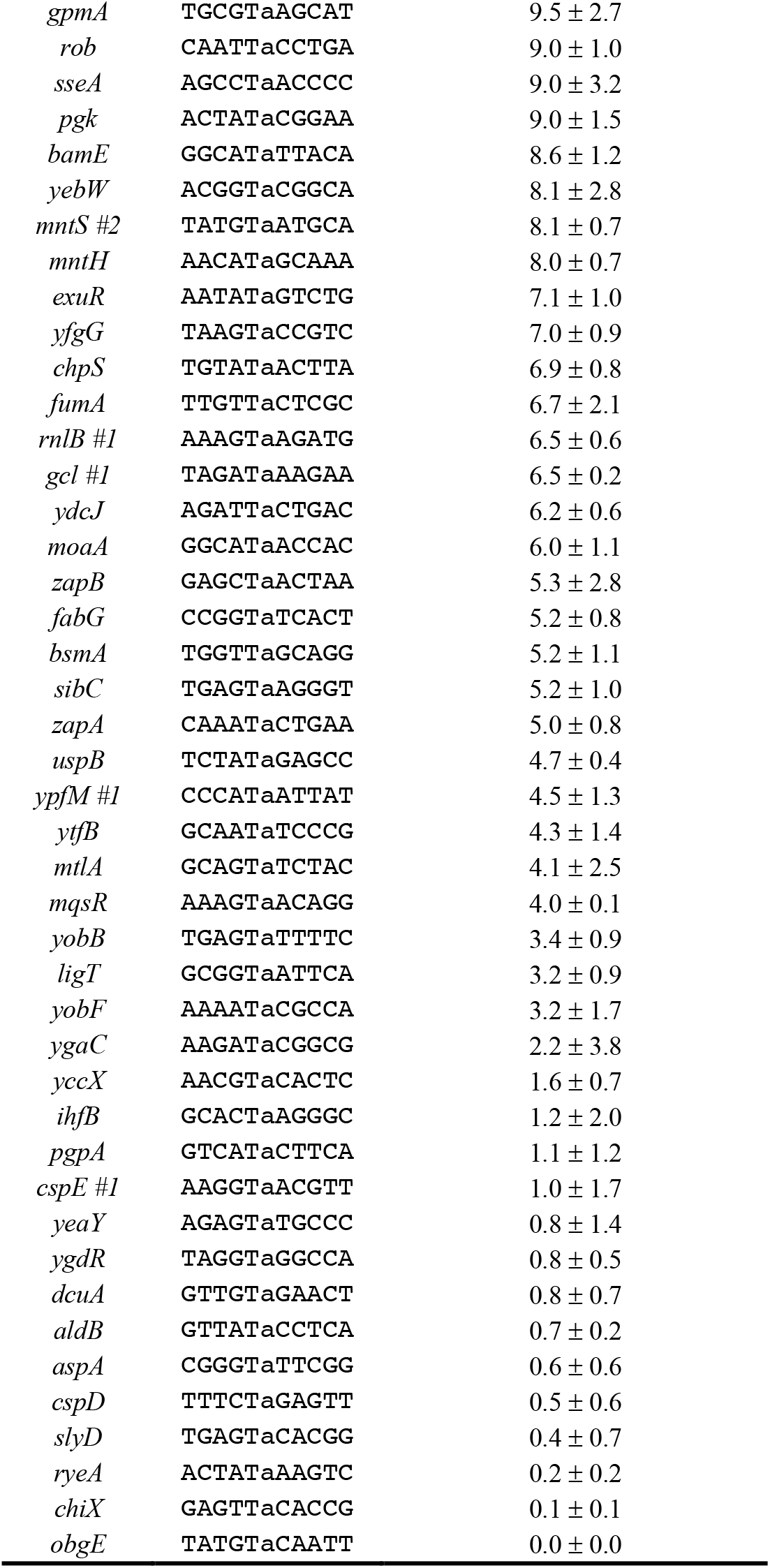
Primer-dependent initiation in natural, chromosomally-encoded *E. coli* promoters.

**Table S2.**
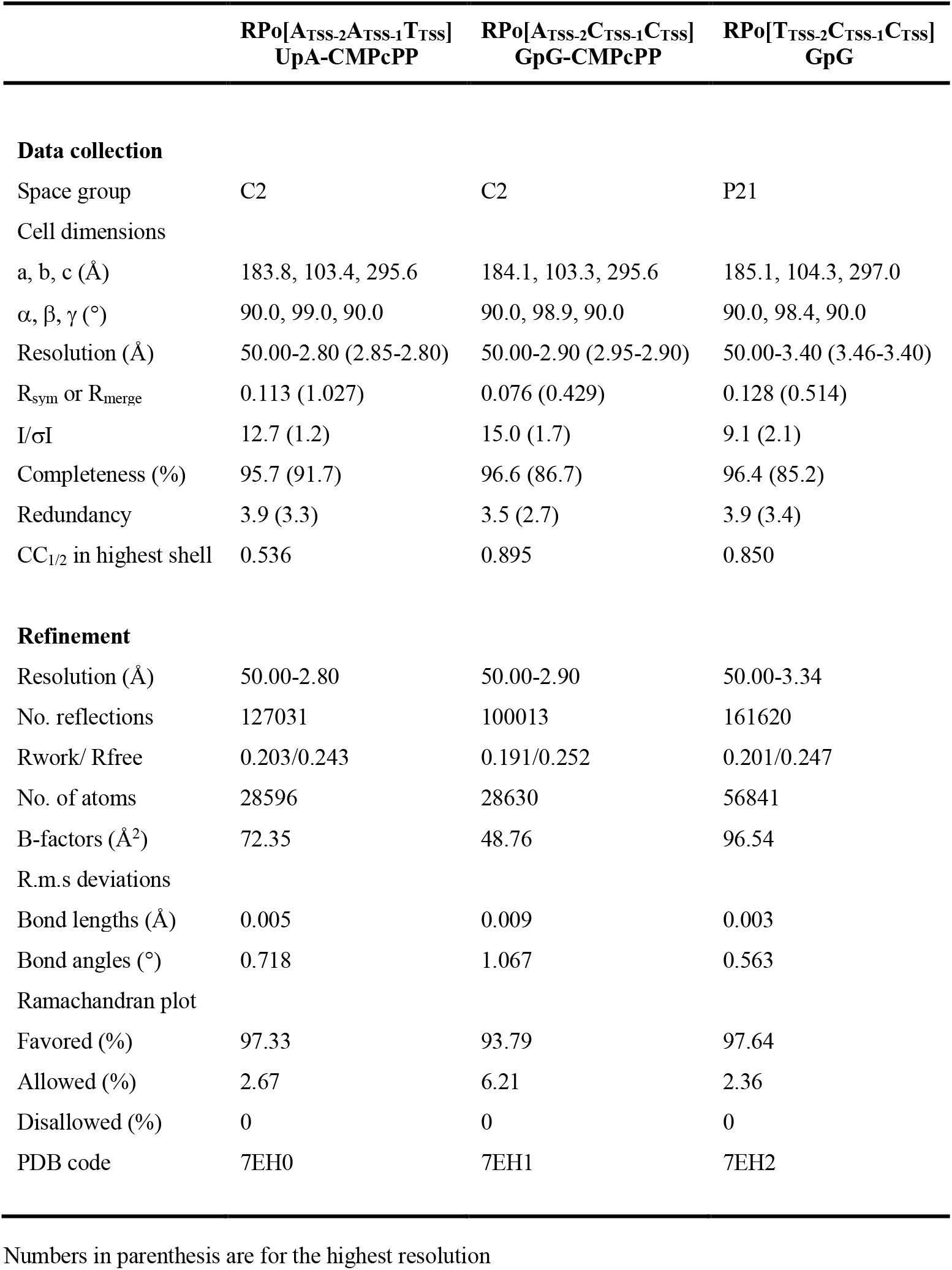
Crystal structure statistics.

**Table S3.**
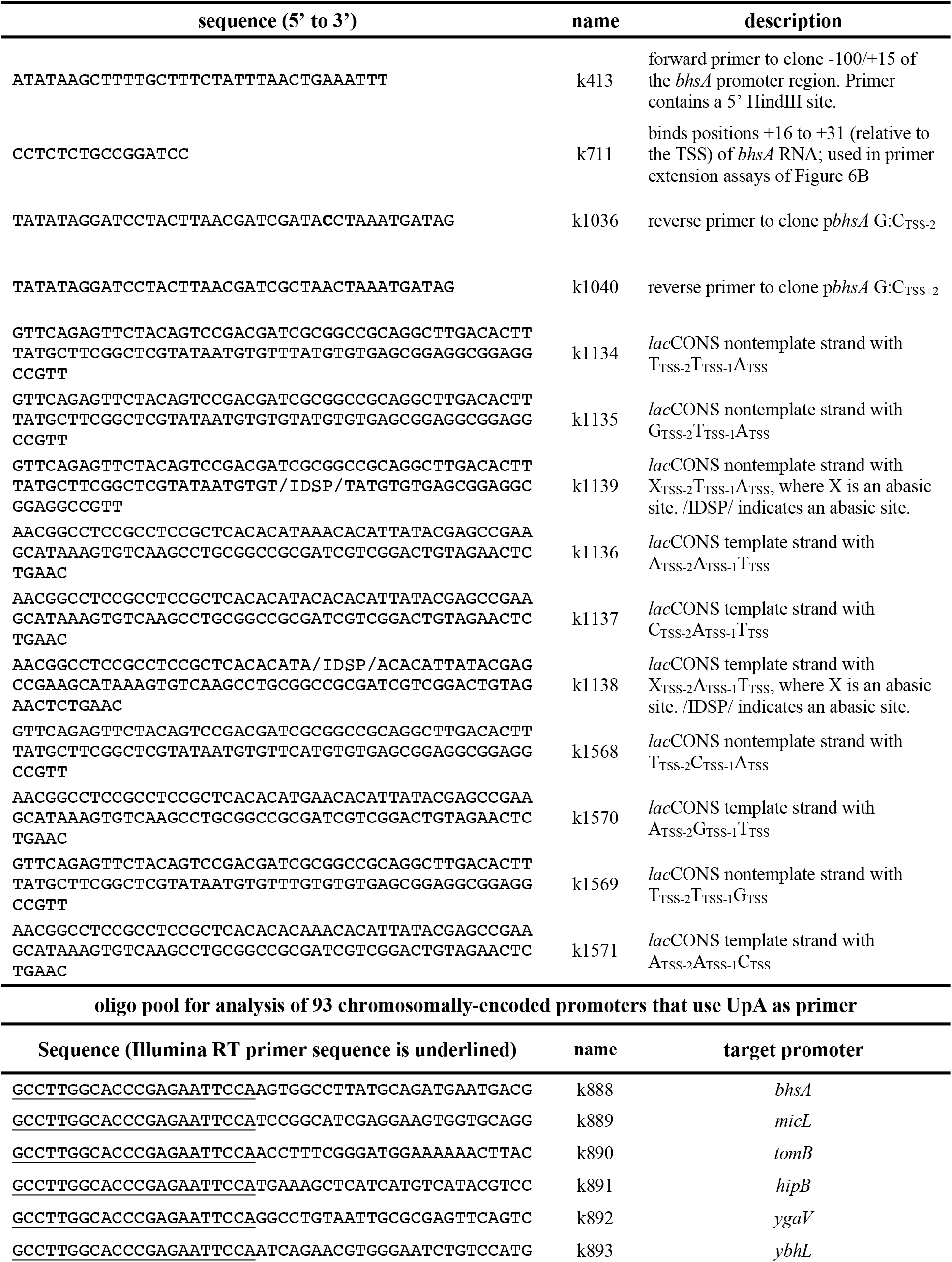
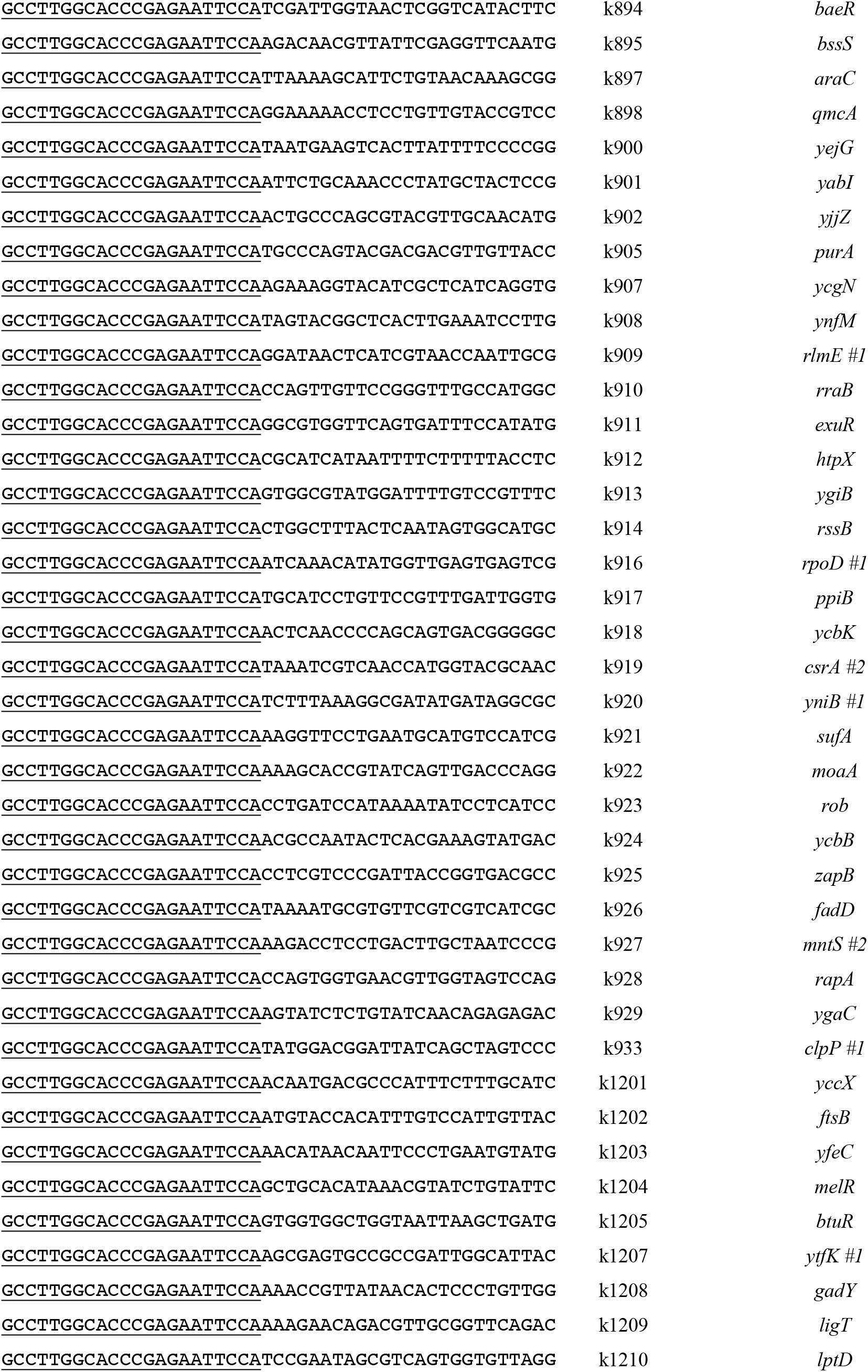
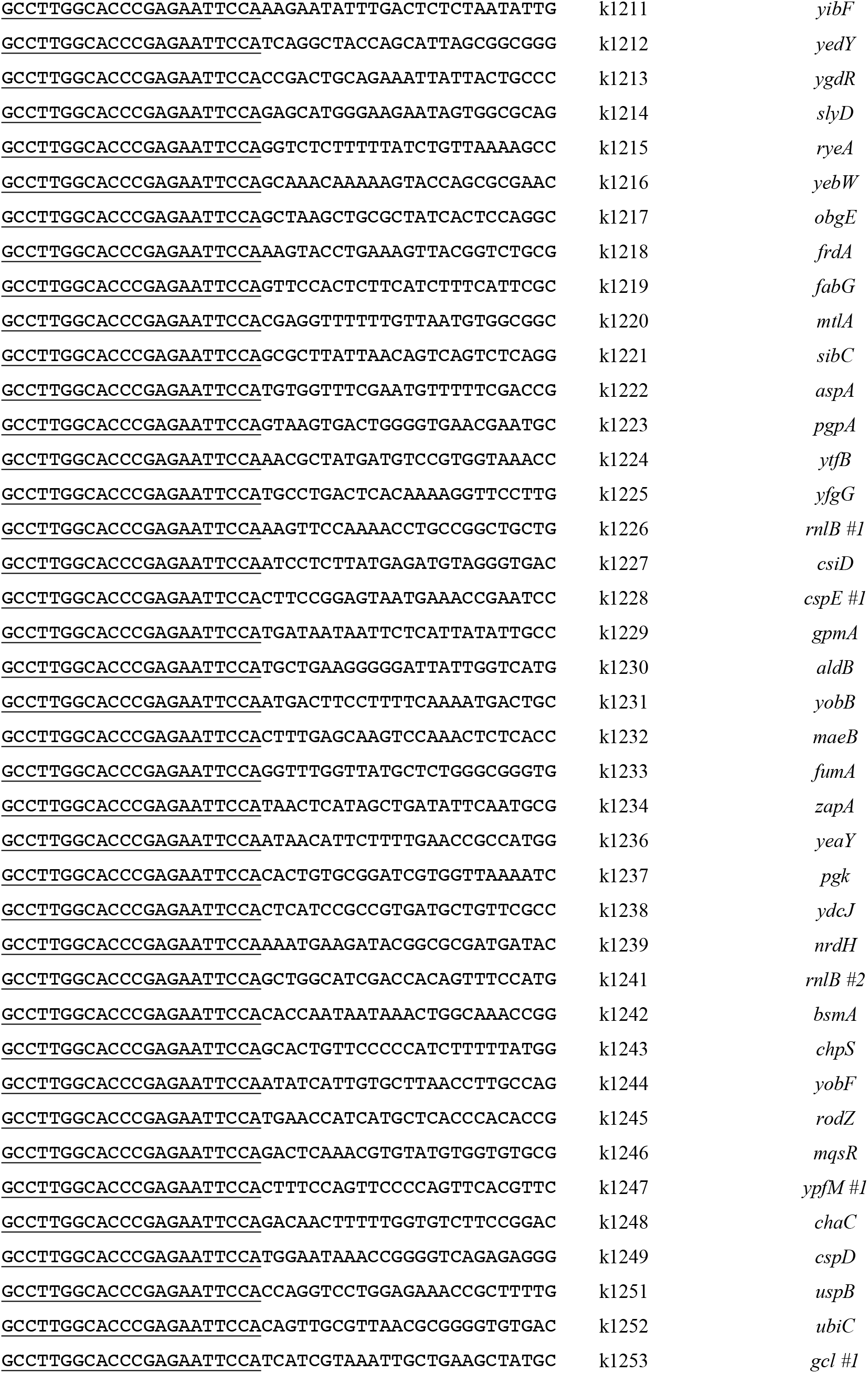
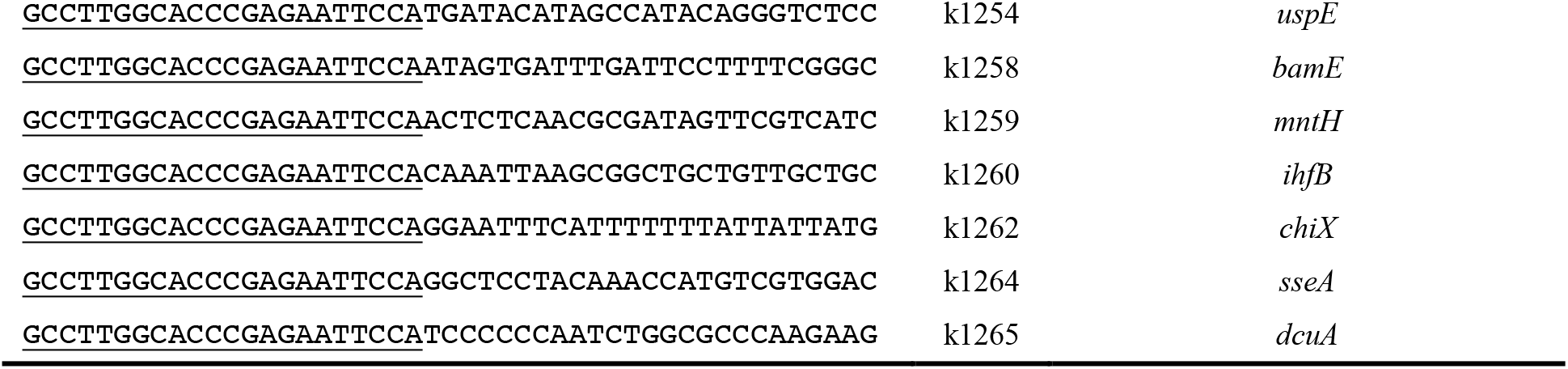
Oligonucleotides.

